# Transcriptional variability accelerates pre-leukemia by cell diversification and perturbation of protein synthesis

**DOI:** 10.1101/2021.10.18.464824

**Authors:** Shikha Gupta, Oliver M. Dovey, Ana Filipa Domingues, Oliwia W. Cyran, Caitlin M. Cash, George Giotopoulos, Justyna Rak, Jonathan Cooper, Malgorzata Gozdecka, Ryan J. Asby, Noor Al-Jabery, Victor Hernandez-Hernandez, Sudhakaran Prabakaran, Brian J. Huntly, George S. Vassiliou, Cristina Pina

## Abstract

Transcriptional variability facilitates stochastic cell diversification and can in turn underpin adaptation to stress or injury. We hypothesize that it may analogously facilitate progression of pre-malignancy to cancer. To investigate this, we initiated pre-leukemia in mouse cells with enhanced transcriptional variability due to conditional disruption of the histone lysine acetyltransferase gene *Kat2a*. By combining single-cell RNA-sequencing of pre-leukemia with functional analysis of transformation, we show that *Kat2a* loss results in global variegation of cell identity and accumulation of pre-leukemic cells. Leukemia progression is subsequently facilitated by destabilization of ribosome biogenesis and protein synthesis, which confer a transient transformation advantage. The contribution of transcriptional variability to early cancer evolution reflects a generic role in promoting cell fate transitions, which, in the case of well-adapted malignancies, contrastingly differentiates and depletes cancer stem cells. In other words, transcriptional variability confers forward momentum to cell fate systems, with differential multi-stage impact throughout cancer evolution.

**One-sentence summary:** Loss of Kat2a enhances transcriptional variability of ribosome biosynthetic programs and transiently accelerates pre-leukemia

---

Tumors evolve by genetic drift and natural selection ^1,2^. Acquisition of new mutations confers a probability of adaptation to new environmental pressures ^3^, and facilitates progression and transformation of pre-malignant lesions, promotes metastasis and drives treatment resistance ^4^. In recent years, it became apparent that non-genetic instability, in particular variability in methylation epialleles, can confer adaptive advantages to tumor growth and survival irrespective of mutations, and function as drivers of therapy resistance and disease relapse in hematological malignancies ^5,6^. Hematological malignancies, and in particular Acute Myeloid Leukemia (AML), are strongly dependent on epigenetic regulation, both through mutation of chromatin factors, and by co-option of unmutated chromatin regulators into maintenance of leukemogenic programs ^7–9^. Notably, AML has lower levels of mutations than solid tumors, supporting the notion that non-genetic events may be especially important in the former ^7^. Akin to genetic instability, epigenetic variability is increased in leukemia initiation and relapse, but low in leukemia maintenance ^10,11^, suggesting that reconfiguration of molecular/transcriptional programs may perturb the identity or survival of well-adapted leukemia cells by disrupting pro-oncogenic molecular signatures. We have recently captured this phenomenon upon loss of KAT2A, a histone acetyltransferase that promotes gene transcription through activation of promoter bursting and stabilization of gene expression levels. *Kat2a* loss (NULL) results in enhanced cell-to-cell transcriptional variability and progressive loss of leukemia stem cells (LSC) transformed with the *KMT2A-MLLT3* (*MLL-AF9*) gene fusion ^12^. Accordingly, KAT2A is required for maintenance of AML cell lines and *in vitro* self-renewal of patient AML blasts ^13^. At a cellular level, loss of *Kat2a* results in perturbation of leukemia lineage trajectories, with emergence of multiple incongruent differentiation pathways that deplete LSC, but fail to uniformly differentiate leukemia cells ^12^. A similar pattern of incongruous exit from the stem cell state was observed upon KAT2A inhibition in mouse embryonic stem (ES) cells ^14^. MLL-AF9 results in an aggressive leukemia, both in mice and in humans, and requires minimal cooperativity from additional mutational events ^7,15^. As such, it provides a good representation of a well-adapted leukemia, with minimal genetic and epigenetic variability. However, it does not reflect what is observed with more common forms of AML such as those associated with *RUNX1-RUNX1T1* (*AML1-ETO*), where progression in mouse models is slow and infrequent ^7,16^, or clonal hematopoiesis, in which the associated mutations (e.g. in *IDH1/2, TET2, DNMT3A*) convey a self-renewal advantage, but require additional genetic events for leukemia ^7,16^. In these cases, we postulate that malignant progression may be facilitated by non-genetic instability, which can be promoted through loss of *Kat2a*.

To test our hypothesis, we made use of 2 pre-leukemia mouse models: *Idh1*^*R132H*^ and *RUNX1-RUNX1T1(RT1(9a))*. First, we developed a new inducible *Idh1*^*R132H*^ allele (Fig. S1a-c, and Supplementary Methods), and crossed it into an *Mx1-Cre* background (Fig. S1d), to activate the mutation in hematopoietic tissues. We verified the functionality of the *Idh1*^*R132H*^ allele by accumulation of the onco-metabolite 2-HG (Fig. S1e-f). *Idh1*^R132H^ mice develop leukemia rarely, with long latency and low penetrance, with no significant effects on overall survival (Fig. S1g). In contrast, combination of *Idh1*^*R132H*^ with other leukemogenic mutations, namely *NRas* and *Npm1c* (triple-mutant), results in short-latency high-penetrance leukemia development (Fig. S1g), confirming the pre-leukemic nature of the *Idh1*^*R132H*^ model. Accordingly, triple-mutant BM cells, but not cells with *Idh1*^*R132H*^ alone, have enhanced colony-forming cell (CFC) assay replating ability, an *in vitro* measure of transformation (Fig. S1h). Comparison of RNA-sequencing from triple-mutant leukemias vs triple-mutant pre-leukemias, or vs *Idh1*^*R132H*^ alone, revealed a gene signature which was specific to the leukemia state, and in which down-regulated genes were enriched for *Kat2a* chromatin targets (Fig. S1i). This association suggests that loss of Kat2a activity may contribute to progression of pre-leukemia to overt AML.

To investigate this putative contribution of *Kat2a* loss to pre-leukemia progression, we crossed conditional *Idh1*^*R132H*^ and *Kat2a*^*Flox/Flox*^ mice, into the Mx1-Cre background (Fig.1a), to generate *Idh1*^*R132H*^ animals that were heterozygous (HET) or NULL for *Kat2a* (Fig. S2a-b). We analyzed *Idh1*^*R132H*^ *Kat2a*^*Flox/WT*^ (*Idh*^*mut*^ *Kat2aHET*) and *Idh1*^*R132H*^ *Kat2a*^*Flox/Flox*^ (*Idh*^*mut*^*Kat2aNULL*) animals 4 and 20 weeks after Cre induction, to identify early and progressed *Idh1*^*R132H*^ pre-leukemia states. Analysis of BM stem and progenitor composition revealed no differences between genotypes or timepoints (Fig.S2c-g). We did not observe differences in spleen or liver pre-leukemia burden (Fig.S2h-i). However, *Idh*^*mut*^ *Kat2aNULL* samples had a significant advantage in CFC re-plating in early pre-leukemia (4 weeks) (Fig.1b), which was not sustained at the 20-week timepoint.

**Fig. 1:**
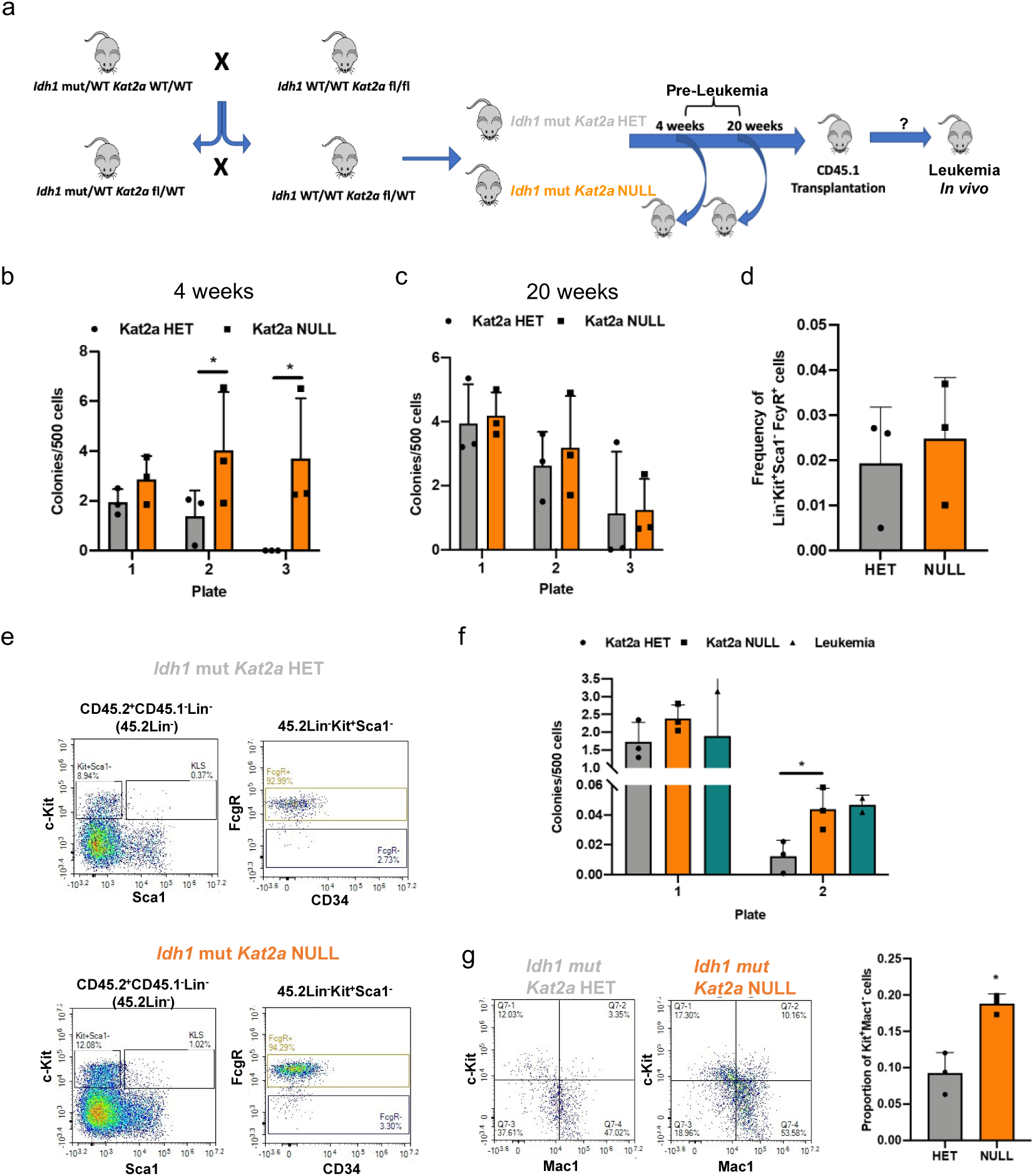
*Kat2a* loss facilitates development of *Idh1*^*R132H*^ pre-leukemia. **(A)** Diagram of *Idh1*^*R132H*^ (*Idh1* mut) and *Kat2a* ^fl/fl^ mouse crosses to generate *Idh1* mut *Kat2a* HET and *Idh1* mut *Kat2a* NULL cells used in pre-leukemia studies. **(B)** Colony-forming cell (CFC) assays of *Idh1* mut *Kat2a* HET and NULL BM cells 4 weeks post-pIpC treatment; mean ± SD, n=3. **(C)** Colony-forming cell (CFC) assays of *Idh1* mut *Kat2a* HET and NULL BM cells 20 weeks post-pIpC treatment; mean ± SD, n=3. **(D)** Quantification of GMP-like BM cells obtained from *Idh1* mut CD45.2^+^ grafts; mean ± SD, n=3 irradiated recipients (CD45.1). **(E)** Representative flow cytometry plots of BM cells in (D); top: *Idh1* mut *Kat2a* HET, bottom: *Idh1* mut *Kat2a* NULL. **(F)** Serial re-plating CFC assays of *Idh1* mut BM grafts. Mean ± SD, n=3 *Idh1* mut *Kat2a* HET and NULL, n=2 *Idh1* mut leukemia. **(G)** Flow cytometry of colonies in (F), Left, representative plots; right, Kit^+^ Mac1^-^ progenitor quantification. Mean ± SD, n=3. All analyses 2-tailed t-test; significant *p<0.05.

This could be compatible with earlier selection of pre-leukemia cells upon *Kat2a* loss, which is achieved later in *Idh*^*mut*^ *Kat2aHET* animals as the *Idh1*^*mut*^ phenotype progresses (Fig. 1c). In an attempt to understand whether the early replating advantage *in vitro* could lead to accelerated leukemia development *in vivo* in the absence of other genetic events, we transplanted BM cells from *Idh*^*mut*^ *Kat2aHET* and *Idh*^*mut*^ *Kat2aNULL* mice, into irradiated CD45.1 recipients and followed them up for 1 year. Similar to single *Idh1*^*mut*^ animals, we could not detect signs of leukemia development in transplanted mice (Fig. S3a). Transplants showed accumulation of GMP-like (Lin-Kit^+^Sca1^-^FcγR^+^) donor cells, compatible with myeloproliferation (Fig. 1d-e), which was identical between genotypes. Peripheral blood counts (Fig.S3b-d) and spleen and liver weights (Fig.S3e-f) were also similar. However, we observed infiltration of spleen and liver in 1 of 3 *Idh*^*mut*^ *Kat2aNULL* recipients, which was not present in *Idh*^*mut*^ *Kat2aHET* grafts (Fig.S3g). Notably, *Idh*^*mut*^ *Kat2aNULL* cells showed enhanced colony-replating potential relative to *Idh*^*mut*^ *Kat2aHET*, which was comparable to that of BM from rare *Idh*^*mut*^ leukemic animals (Fig. 1f). *Idh*^*mut*^ *Kat2aNULL* cells in CFC assays were enriched in c-Kit^+^Mac1^-^ cells (Fig. 1g) compatible with hindered differentiation and/or expansion of self-renewing cells. Overall, the results suggest that loss of *Kat2a* imparts leukemogenic properties to *Idh1*^*mut*^ cells but is in itself not sufficient to drive leukemogenesis in the absence of additional cooperating genetic events.

We next tested the impact of *Kat2a* loss on the pre-leukemia model driven by the exon 9a splicing variant of the *RUNX1-RUNX1T1* (*RT1(9a)*) fusion gene, which when retrovirally-delivered to adult BM cells, leads to long-latency, incomplete-penetrance leukemia in irradiated recipients. Using our previously described *Kat2a*^*Flox/Flox*^ *Mx1-Cre* mice, we isolated progenitor-enriched BM cells after *pIpC*-induced locus excision (Fig. S4a), and delivered the RT1(9a) construct by retroviral transduction, as described ^17^. In all experiments, *Kat2a*^*Flox/Flox*^ *Mx1-Cre*^*+/-*^ (*Kat2aNULL*) were compared with *Kat2a*^*Flox/Flox*^ *Mx1-Cre*^*-/-*^ (*Kat2aWT*) cells. We started by evaluating leukemia development after transplantation of RT1(9a) *Kat2aNULL* and *Kat2aWT* BM cells (Fig. 2a). Loss of *Kat2a* led to a dramatic decrease in survival of RT1(9a) recipient animals, compatible with accelerated leukemia progression (Fig. 2b). *Kat2a NULL* leukemias had a non-significant trend towards higher white blood cell counts (Fig. S4b-d) and spleen leukemia burden, with minimal infiltration of other organs (Fig. S4e-g). The surface phenotype of the leukemias was indistinguishable between genotypes (Fig. S4h). Analysis of early timepoints post-transplantation showed that RT1(9a) engraftment became quickly fixed in the absence of *Kat2a* (Fig. 2c). *Kat2aNULL*/*RT1(9a)* cells obtained from healthy pre-symptomatic recipients were enriched for GMP-like cells (Fig. 2d), and displayed enhanced colony formation (Fig. 2e), compatible with accelerated pre-leukemia development. Similarly, *Kat2aNULL* cells directly tested in CFC assays upon retroviral transduction, displayed enhanced re-plating potential. (Fig. 2f). In contrast, excision of *Kat2a* in RT1(9a) cells post-*in vitro* transformation by 3 rounds of serial-replating, led to a reduction in colony formation (Fig. 2g), suggesting that *Kat2a* loss favors leukemia development only at a pre-leukemia stage. These observations mirror our previously identified role for *Kat2a* in maintenance of established leukemia stem-like cells and suggest that *Kat2a* plays stage-specific roles during leukemogenesis, which are preserved across leukemia models.

**Fig. 2:**
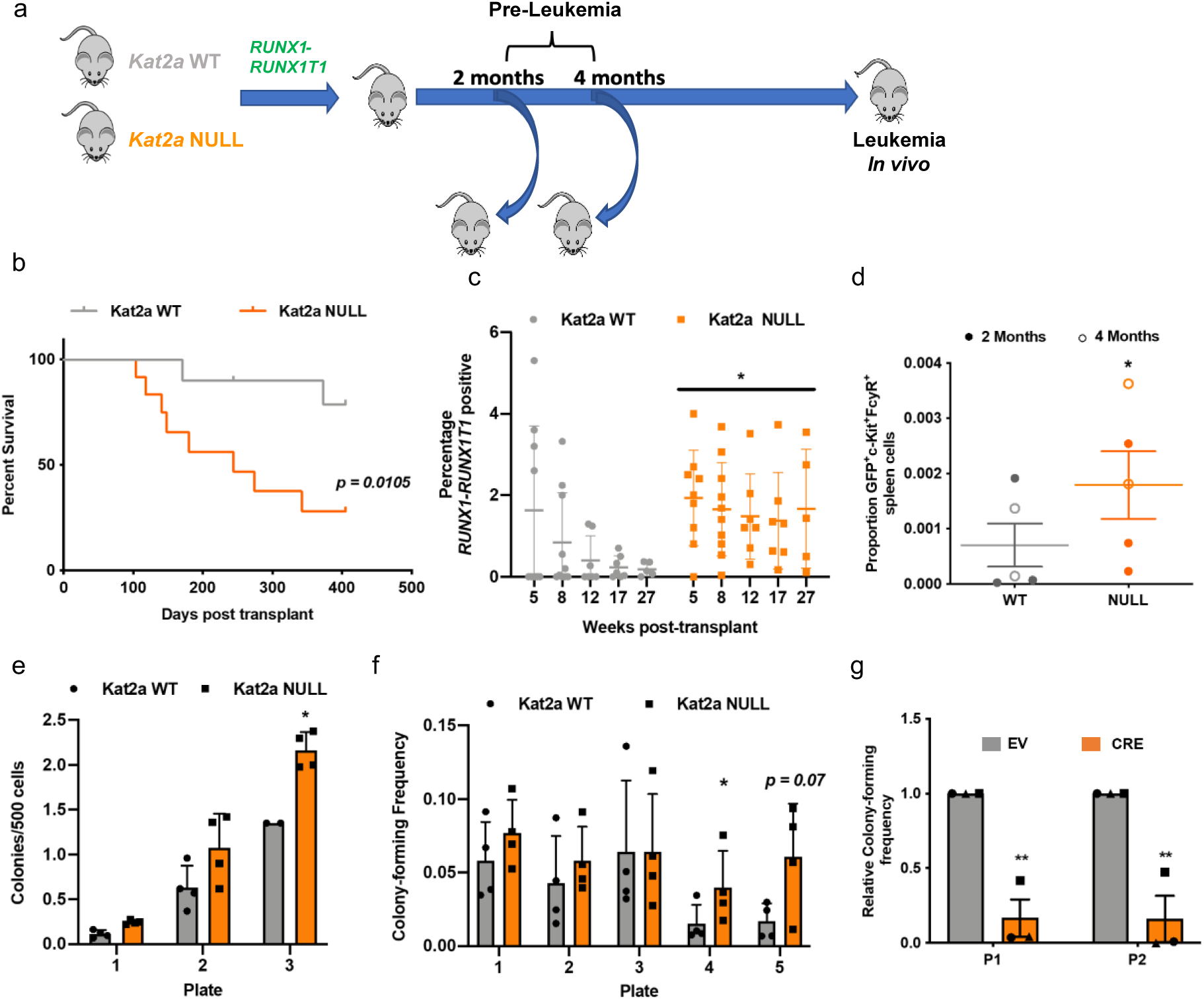
*Kat2a* loss accelerates *RT1(9a)* pre-leukemia to leukemia progression. **(A)** Experimental design. **(B)** Survival curve of *RT1(9a) Kat2aWT* and *Kat2aNULL* Kit^+^ BM recipients; n=12 animals/genotype, *p<0.05, log-rank test. **(C)** Quantification of peripheral blood GFP for animals in (A); GFP reports *RT1(9a)*. Mean ± SD, n=10 animals/genotype (8 weeks), *p<0.05, 2-way ANOVA. **(D)** CFC assay of *RT1(9a) Kat2aWT* and *Kat2aNULL* graft BM cells 4 months post-transplantation; mean ± SD, n=4. **(E)** Flow cytometry analysis of *RT1(9a) Kat2aWT* and *Kat2aNULL* graft spleen cells 2- and 4-months post-transplantation; mean ± SD, n=5. **(F)** *In vitro* transformation of *Kat2aWT* and *Kat2aNULL* Lin^-^/Kit^+^ BM cells transduced with *RT1(9a)* retrovirus tested in CFC serial re-plating; mean ± SD, n=4. **(G)** CFC re-plating (plate=P1, P2) analysis of *RT1(9a) Kat2a*^*Flox/Flox*^ Cre^-/-^ Kit^+^/Lin^-^ BM cells excised *in vitro* by lentiviral-delivered *Cre* recombinase (vs. EV, empty vector) after 3 rounds of colony re-plating. Mean ± SD, n=3. All other analyses 2-tailed t-test, *p<0.05, **p<0.01.

We had previously associated *Kat2a* function in leukemia stem cell maintenance with stability of transcriptional programs ^12^. Using single-cell RNA-sequencing (scRNA-seq), we showed that *Kat2a* loss resulted in diversification and branching of differentiation trajectories, and associated with enhanced transcriptional noise, particularly in biosynthetic programs (e.g., ribosomal biogenesis and translation). We asked if similar mechanisms were at play in pre-leukemia progression facilitated by *Kat2a* loss. We hypothesized that enhanced transcriptional variability leading to program diversification might increase the probability of accessing or seeding leukemia programs, resulting in the observed acceleration in leukemia progression. We performed scRNA-seq analysis of pre-leukemia cells on the 10X platform, comparing transcriptional landscapes of *Kat2a*NULL and *Kat2a*WT RT1(9a) asymptomatic animals obtained 2 and 4 month post-transplantation. We sequenced a total of 1767 cells sorted as RT1(9a)/GFP^+^ Kit^+^ stem/progenitor and retrieved an average of 174770 aligned reads per cell, corresponding to medians of 5939 Unique Molecular Identifiers (UMI) and 1575 genes per cell (Supplementary File 1). Less than 0.2% of reads aligned to mitochondrial DNA, denoting successful sequencing. Pre-processing steps are detailed in Supplementary Methods.

We employed transcripts of cell surface markers routinely used for hematopoietic cell immunophenotyping to map the identity of cells along the pseudo-temporal trajectories (Fig. S5a-b). Cells at the origin of the trajectory expressed high *Ly6e* (*Sca1*), *Cd34* and *Flt3*, compatible with lymphoid-myeloid-primed progenitors (LMPP). LMPPs were adjacent to a granulocyte-monocyte progenitor (GMP)-like state (*Ly6e*^low^*Cd34*^+^*Fcgr3*^+^). Trajectories involved 3 additional states: *Ly6e*^+^ *CD79a*^+^ *Cd14*^-^ B-cell affiliated Progenitor (BAP) (Supplementary File 3), and *Ly6e*^+^*Fcgr3*^+^*Cd14*^+^ Monocyte-affiliated Progenitor (MAP) (Supplementary File 4), confirmed by gene ontology (GO) analysis (Fig. S5c-d); and a third compartment in direct proximity of the GMP, characterized as *Ly6e*^+^*Fcgr3*^+^*Cd33*^+^*Cd14*^*+*^, with no*Cd34* or *Cd48*. BAP was exclusive to *Kat2a*NULL samples, while MAP was common to both genotypes, albeit enriched in *Kat2a*NULL samples (Fig. S5e). We confirmed enhanced phenotypic differentiation of *Kat2a*NULL RT1(9a) cells to the B (Fig. 3c) and macrophage (Fig. 3d) lineages *in vitro*. Using a signature of RT1 chromatin targets ^18^, we identified the third compartment as the candidate pre-leukemia progenitor (PLP) population (Fig. 3e-f). PLPs form a discrete (Fig. 3e) and relatively smaller (Fig. S5e) compartment in the *Kat2a*WT trajectory; in contrast, the GMP-to-PLP transition is more densely populated in *Kat2a*NULL pre-leukemia (Fig. 3f), with PLP comprising a larger number of NULL cells. (Fig. S5e). Overall, the pseudo-temporal trajectories support the notion of increased cell diversification through *Kat2a* loss, with additional cell states (BAP) and, importantly, increased size of the PLP compartment. Comparative analyses of 2 and 4-month pre-leukemia samples confirm the pseudo-temporal trajectory findings: *Kat2a*WT RT1(9a) cells progressively differentiate from LMPP to GMP-like cells (Fig. S5f) and accumulate PLPs. Pre-leukemia progression is accelerated in *Kat2a*NULL RT1(9a) (Fig. S5f), which contain nearly 50% of PLPs at 4 months, and uniquely display 27% of BAP cells at 2 months (Fig. S5f). The increased diversification of cell types in *Kat2a* NULL samples translated in enhanced cell-to-cell transcriptional variability, measured by pairwise distance ^19^ (Fig. 3g, S6a). *Kat2a*NULL transcriptional programs were also more variable within individual cell sub-populations (Fig. S6b), indicating that the variability is likely to at least in part reflect transcriptional noise, rather than differences in cellular composition alone. Comparison of transcriptional variability between *Kat2a* WT and NULL cell subpopulations shows that the GMP to PLP transition itself is accompanied by enhanced transcriptional variability (Fig. S6b), supporting the notion that *Kat2a* loss may accelerate pre-leukemia progression through enhanced transcriptional noise.

**Fig. 3:**
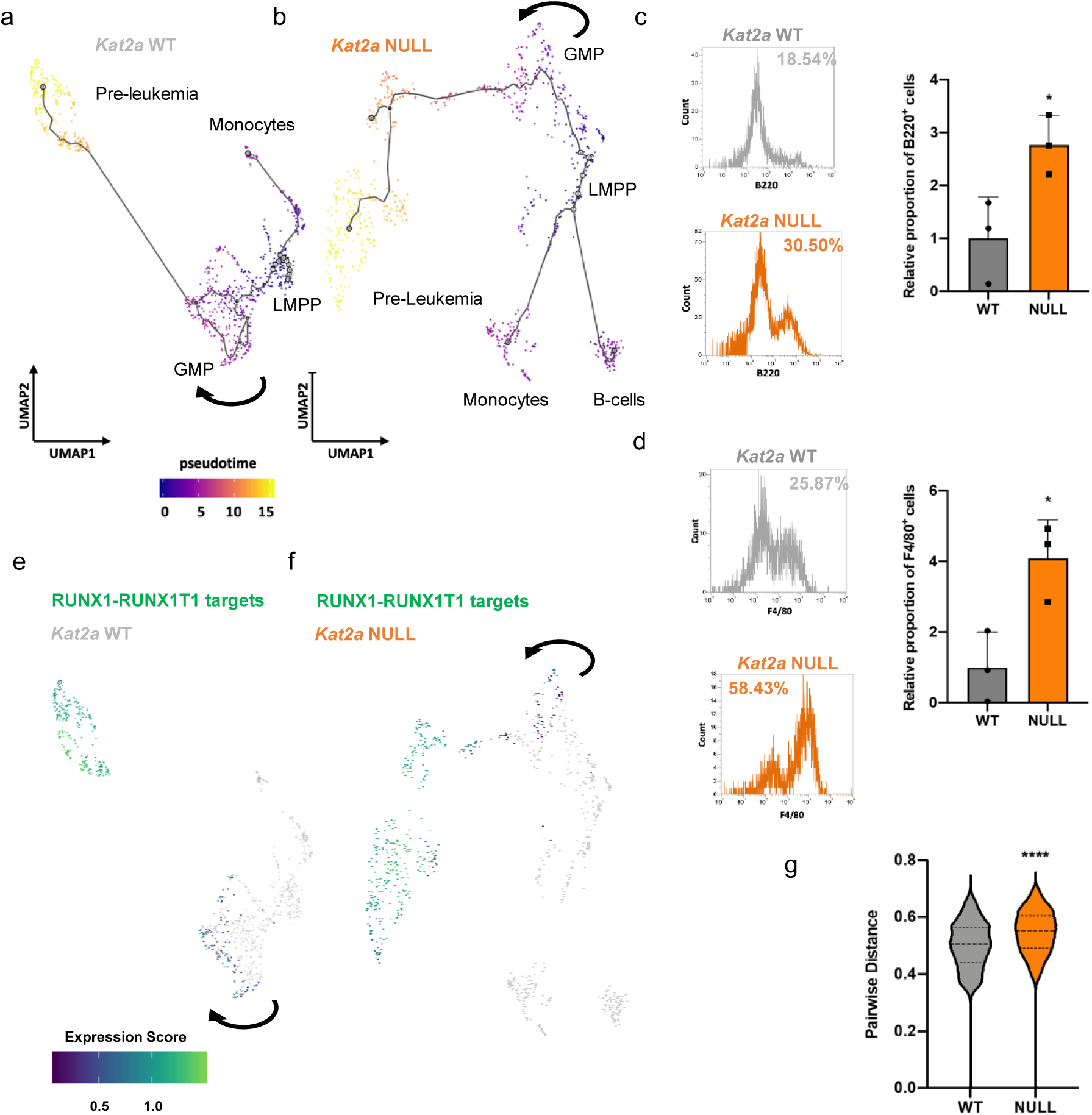
Loss of *Kat2a* diversifies cell fates and promotes RT1(9a) pre-leukemia progression. **(A-B)** Pseudotime single-cell trajectory of **(A)** *Kat2aWT* cells, **(B)** *Kat2aNULL* RT1(9a) cells 2 and 4 months after transplantation. Trajectories inferred using Monocle3 ^21^; compartments labelled as per hematopoietic markers in Fig. S5A-B. Arrows denote pseudotime progression. **(C)** B220 B-cell marker in plate 2 CFC of RT1(9a)-transduced *Kat2aWT* and *Kat2aNULL* cells during *in vitro* transformation; mean ± SD, n=3. **(D)** F4/80 monocyte marker in plate 2 CFC of RT1(9a)-transduced *Kat2aWT* and *Kat2aNULL* cells during *in vitro* transformation; mean ± SD, n=3. **(E-F)** Expression of *RUNX1-RUNX1T1* ChIP-seq targets ^18^ in **(E)** *Kat2aWT* cells and **(F)** *Kat2aNULL* RT1(9a) single-cell trajectories. Arrows as in A-B. **(G)** Pairwise distance transcriptional variability measure^19^ of *Kat2aWT* and *Kat2aNULL* RT1(9a) cells; top 500 most variable genes/genotype calculated by distance to the median CV (DM); ****p-adj<0.0001. All analyses 2-tailed t-test, *p<0.05.

In order to understand the nature of the transcriptional programs perturbed upon (1) *Kat2a* loss, and (2) pre-leukemia progression, we performed differential gene expression analysis of the scRNA-seq dataset. Comparison of *Kat2a*NULL to WT cells revealed minimal changes in gene expression levels (Fig. S7a), which were of down-regulation, as previously observed upon *Kat2a* loss ^12^. Consistent with our published data ^12^, differentially-expressed genes between genotypes predominantly associated with ribosomal assembly and translation ontologies (Fig. S7b) (Supplementary File 5), a pattern particularly prominent within PLP (Fig. 4a) (Supplementary File 6). The same ontologies were specifically down-regulated in *Kat2a*WT RT1(9a) PLPs compared to other cell states (Fig. S7c) (Supplementary File 7), capturing a reported decrease in protein synthesis in RT1 leukemia ^20^. Ribosomal and translation ontologies (Fig. S7d-e) (Supplementary File 8) were also down-regulated in *Idh1*^*R132H*^ mice upon pre-leukemia-to-AML progression through additional genetic mutations. Altogether, our findings suggest a specific association of attenuated ribosomal programs with pre-leukemia progression, which may be further facilitated by *Kat2a* loss. *Kat2a* loss increases variability of ribosomal biogenesis programs in PLPs (Fig. 4b), themselves more variable than GMPs (Fig. S7f), suggesting enhanced noise at the transition (Supplementary File 9). The gene expression range in *Kat2a*NULL PLPs favors lower mean values (Fig. 4c). In support of the functional impact of the transcriptional perturbation, *Kat2a* loss results in decreased protein synthesis (Fig. S8a-b; also ^12^).

**Fig. 4:**
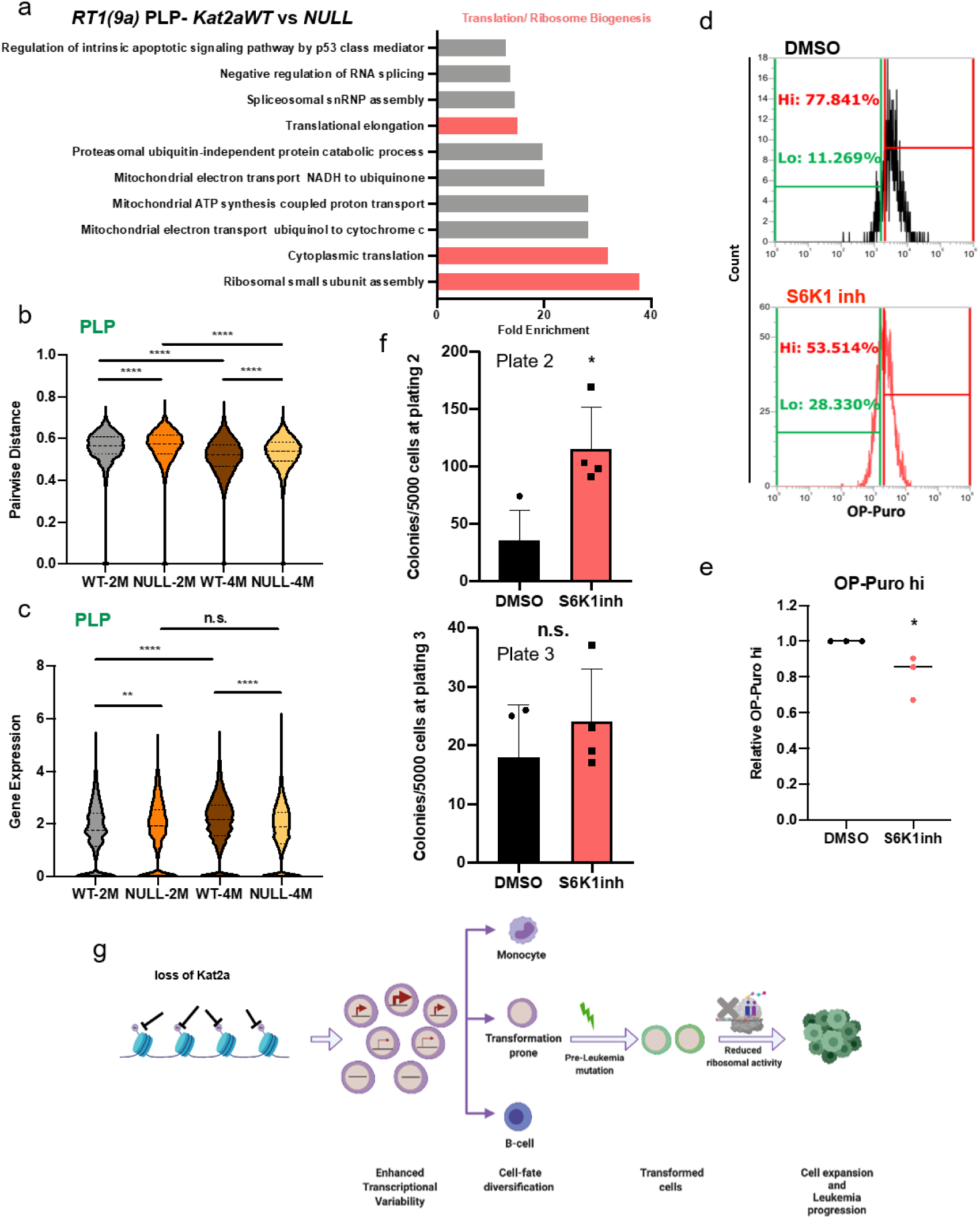
Inhibition of protein synthesis phenocopies effects of *Kat2a* loss facilitating pre-leukemia transformation. **(A)** Over-represented gene ontology categories (GO) for genes downregulated in RT1(9a) *Kat2aNULL* vs. *WT* PLP; *p-adj<0.05. **(B)** Pairwise distance ^19^ of RT1(9a) PLPs; comparisons consider correlations between ribosomal biogenesis genes ****p-adj<0.0001, 2-tailed t-test. **(C)** Distribution of expression levels for gene signatures in (B) ****p-adj<0.0001, 2-tailed t-test. **(D)** Representative OP-Puro incorporation flow cytometry of S6K1inh-treated RT1(9a) *Kat2aWT* cells. **(E)** Quantification of OP-Puro high cells in (E), relative to DMSO; mean ± SD, n=3, *p<0.05, 2-tailed t-test. **(F)** CFC replating of *Kat2aWT* RT1(9a) *in vitro* transformation in the presence of S6K1inh (control, DMSO). Plate 2 (left); mean ± SD, n=4, *p<0.05. Plate 3 (right); mean ± SD, n=4, n.s; 2-tailed t-test. **(G)** Proposed mode of action of *Kat2a* loss in pre-leukemia progression.

We tested the contribution of reduced protein synthesis activity to pre-leukemia progression by treatment with the S6K1 inhibitor (S6K1inh) PF4708671 (Fig. S8c), which impairs protein synthesis activity confirmed by reduced OP-Puro incorporation in nascent peptide chains (Fig. 4d-e). We treated *Kat2a*WT RT1(9a) cells with S6K1inh and tested their leukemia transformation potential *in vitro* through CFC assay re-plating. S6K1-inhibited cells displayed enhanced colony-formation upon re-plating (Fig. 4f), suggesting a contribution to leukemia transformation. However, the increase in colony formation was transient and eventually lost upon subsequent re-plating (Fig. 4f). This suggests that the effects of reduced protein synthesis on leukemia cells may vary with progression of transformation, reconciling our data with prior analysis of established *MLL-AF9* cells, in which reduced OP-Puro incorporation associated with *Kat2a*NULL-mediated extinction of leukemia stem cells ^12^. We observed a similar pattern of transient increase in colony formation of *Idh1*^*R132H*^ pre-leukemia cells treated with S6K1inh (Fig. S8d). Altogether, the data suggest that reduced ribosomal assembly and protein synthesis facilitate pre-leukemia progression. Exploration of lower levels of expression of translation-associated genes as a consequence of enhanced transcriptional variability may be instrumental in the acceleration of pre-leukemia to AML transition upon *Kat2a* loss. As leukemia progresses, variability in ribosomal biosynthesis programs, may become attenuated with deviation from an optimal level no longer favorable to transformation.

In this report, we have shown that *Kat2a* loss facilitates pre-leukemia progression in *Idh1*^*R132H*^ and *RUNX1-RUNX1T1(9a)* mouse models of human disease, with acceleration of frank leukemia onset in the case of RT1(9a). Loss of *Kat2a* resulted in enhanced variability of transcription, leading to diversification of cell fates, including accumulation of pre-leukemia progenitor cells. In the context of an early genetic event such as *RT1(9a)* or *Idh1*^*R132H*^, which do not allow for full leukemia transformation, the cellular heterogeneity that ensues creates the opportunity for specification and expansion of transformation-prone cells, on which additional molecular events may act to progress the leukemic process (Fig. 4g). Amongst these, we show that destabilization of translation, which is specifically targeted by *Kat2a*, acts to facilitate transformation. This may be achieved by surveying and selection of biosynthetically quiescent cell states, which evade further diversification and respond to additional mutations with disease propagation and progression. Fully transformed, well-adapted leukemia cells may buffer transcriptional variability to maintain stable self-renewal signatures and optimal biosynthetic, translation rates. In this context, instability of transcriptional programs may shift biosynthetic homeostasis and perturb cellular identity, and mal-adapt leukemia stem-like cells, with anti-leukemia effects. Thus, stage-specific tuning and untuning of transcription and translation may be employed to modulate cancer progression, a principle that can be extended to other cancer state transitions such as metastasis or drug-resistance with prognostic and therapeutic potential.

## AUTHOR CONTRIBUTIONS

Study conception – CP; Experimental design – SG, OD, GSV, CP; Data collection – SG, OD, AFD, OC, CC, GG, JR, JC, MG, RJA, NA, VH-H; Data analysis and interpretation – SG, OD, AFD, OC, CC, GSV, CP; Critical reagents – SP, BJH; Writing – SG, CP, with contributions from OD, GSV. All Authors approved the final version of the manuscript.

The Authors would like to thank: Central Biomedical Services of the University of Cambridge for expert animal husbandry, CRUK Genomics Core Facility at the Cambridge Research Institute, and the Wellcome Trust Sanger Research Institute Genomics Core Facility for library preparation and next-generation sequencing; the Flow Cytometry facilities at the Cambridge Institute for Medical Research, the NIHR Cambridge BRC Cell Phenotyping Hub, and the Department of Pathology of the University of Cambridge (Dr Joana Cerveira) for cell sorting; Dr Roberto Bandiera for helpful discussions and reagent sharing; and Dr Matt Wayland for assistance with Cell Ranger installation and 10X Genomics data matrix generation.

## DATA AVAILABILITY

Single-cell RNA sequencing and bulk RNA-sequencing data were deposited in ArrayExpress with accession number E-MTAB-10853 and ERP006862 respectively.

## SUPPLEMENTARY MATERIALS

Materials and Methods

Table S1-S3

Fig S1-S8

References 26-32

Supplementary Files S1-S9

