## Supplementary Methods, Figures and Figure Legends for "Transcriptional variability accelerates pre-leukemia by cell diversification and perturbation of protein synthesis"

### SUPPLEMENTARY MATERIALS

#### SUPPLEMENTARY METHODS

##### Pre-leukemia mouse models

Mice were kept in a Specific Pathogen Free (SPF) animal facility, and all experimental work was carried out under UK Home Office regulations. Animal research was regulated under the Animals (Scientific Procedures) Act 1986 Amendment Regulations 2012 following ethical review by the University of Cambridge Animal Welfare and Ethical Review Body (AWERB). Peripheral blood was collected by saphenous vein and differential blood cells counts were determined using a Vet abc automated counter (Scil Animal Care, Viernheim, Germany).

##### Generation of an *Idh1*$^{R132H}$ mouse model

Targeting vector was generated as follows using methods described previously [22]. The wildtype *Idh1* locus (endogenous *Idh1* sequence, including arms of homology were captured by gap repair as previously described). The following primer pairs were used to amplify the 'U' cassette containing attR1 and attR2 gateway cloning sites containing a *Zeo*/*PheoR* selection cassette with the appropriate overhangs to allow insertion of this cassette between exons 2 and 3 of the *Idh1* locus by recombineering (Table S1)

The following primers were used to amplify the 'G' cassette which contains attR3 and attR4 Gateway sites flanking an *AmpR* cassette. Appropriate overhangs were incorporated into these primers to allow 'Gap repair' subcloning and retrieval of arms of homology 5' and 3' to exons 2 and 4 (5.9 and 3.6kb respectively) from the *Idh1* containing BAC (Table S1).

A custom gene block (GeneArt, ThermoFisher) containing sequence encoding the mutant R132H substitution in exon 3 was cloned into the subsequent U/G captured intermediate - replacing the wildtype exon3 by standard restriction enzyme cloning using S*naBI* and *CspCI* to generate a R132H mutant U/G vector. A custom cDNA flanked by *AflII* and *AscI* sites, containing AttL1-loxP-En2SA-Idh1 cDNA exons3-9-SV40 pA-loxP-FRT was synthesised (GeneArt ,ThermoFisher) and cloned into the PL1PL2 containing a FRT flanked NeoR cassette. This generated the 'SA-Idh1 exon 3-9 cDNA' cassette.

These vectors and the PL3L4 vector were combined in the downstream L/R clonase reaction to successfully generate the Idh1R132H-*NeoR*TV vector. All intermediate and final vectors were sequenced verified.

The *Idh1$^{R132H-NeoRTV}$* targeting vector was electroporated into mouse ES cells and genotyping was performed for on target integration at the endogenous Idh1 locus using a series of long-range PCR reactions (Table S1):

GDNA extracted from heterozygous targeted single cell mouse ES cell clones were subjected to Southern Blot to confirm the structural integrity and confirmation of site directed recombination of FRT and LoxP recombination sites, prior to microinjection. FRT recombination and removal of the *NeoR* cassette was mediated by expression of flippase via transient transfection of the pCAG-FlpO plasmid (addgene #89574). LoxP recombination and deletion of the 'SA-Idh1 exon 3-9 cDNA' cassette was mediated by expression of Cre via transient transfection of the pCAG-Cre plasmid (addgene #13775). GDNA extracted from single cell clones were subjected to Southern Blot to confirm the successful removal of the *NeoR* resistance cassette and LoxP recombination to generate the *Idh1$^{R132H-TV}$* and *Idh1$^{R132H-KI}$* alleles respectively (Fig. S1a). The primers which were used to generate Southern Blot hybridisation probes for the respective 3'internal (FLP assay) and 5'internal (Cre assay) are mentioned in Table S1.

Positively targeted heterozygous clones were selected for microinjection. Chimeric offspring were then selected for downstream breeding and germ line transmission of the *Idh1$^{R132H-NeoRTV}$* targeted allele.

The FRT flanked Neomycin resistant cassette, used for positive enrichment of targeted mouse ES cells, was removed by breeding to FLPe mice (as previously described- [23]). F1 mice were backcrossed to wildtype C57Bl6 mice and mice negative for the presence of the RosaFLPe transgene and positive for the inducible *Idh1Knock-In$^{R132H}$* allele were selected for downstream cohort generation by subsequent crosses with the inducible Mx1-Cre transgenic mouse model (Fig. S1c). Standard PCR genotyping for the wildtype and mutant alleles was performed (using the primers detailed in Table S1). Subsequent crosses to homozygous *Nras$^{G12D/G12D}$* and *Npm1$^{cA/cA}$* cohorts were used to generate experimental model cohorts (as described previously, [24]).

In order to generate a Mx1-Cre-inducible mouse with *Idh1*$^{mut/WT}$ and *Kat2a* floxed alleles, *Idh1*$^{mut/WT}$ *Kat2a*$^{WT/WT}$ *Mx1-Cre*$^{+/-}$ males were crossed with *Idh1*$^{WT/WT}$ *Kat2a*$^{fl/fl}$ *Cre*$^{-/-}$ females. The first-generation carried a *Kat2a*$^{fl/WT}$ genotype (referred as *Kat2a* HET) with either *Idh1*$^{WT/WT}$ or *Idh1*$^{mut/WT}$ and *Mx1-Cre*$^{+/-}$. *Idh1*$^{WT/WT}$ *Kat2a*$^{fl/WT}$ and *Idh1*$^{mut/WT}$ *Kat2a*$^{fl/fl}$ offspring were crossed to obtain experimental genotypes referred to as *Kat2a* HET- *Idh1*$^{mut/WT}$ *Kat2a*$^{fl/WT}$ and *Kat2a* NULL-*Idh1*$^{mut/WT}$ *Kat2a*$^{fl/fl}$ maintaining a heterozygous *Mx1-Cre* allele.

**_Kat2a_ conditional knockout model**

*Kat2a*$^{fl/fl}$ conditional knockout mice have been previously described [12].

**Genotyping**

Ear notch biopsies were digested overnight in lysis buffer (50mM Sodium Chloride, 50mM Tris(hydroxymethyl)aminomethane hydrochloride, 5mM Ethylenediaminetetraacetic acid, 20% Sodium dodecyl sulphate, 0.5mg/ml Proteinase K) at 55°C, 750 rpm using a Thermoshaker (BioSan). DNA extraction used isopropanol-based precipitation. Mice were genotyped using the primers in Table S1, following the PCR protocol: 95°C, 3min, 40X (94°C, 30sec; 60°C, 30 sec [57°C, 30sec for *Idh1*]; 72°C, 90sec [30 sec for *Idh1*]); 72°C, 10min. DNA products were run on a 1% Agarose Gel in TAE (1x), at 100V and visualized using an AlphaImager UV transilluminator (Protein Simple). Cre-mediated recombination was induced in 8-weeks old mice by administration of 5 alternate-day intraperitoneal injections of poly(I)- poly(C) (pIpC), 300 µg/dose. *Idh1* recombination was confirmed by PCR (Table S1, reverse primer 2) following the same PCR protocol.

**Table S1: Primers for genotyping**

| Target | Forward primer (5’-3’) | Reverse primer 1 (5’-3’) | Reverse primer 2 (5’-3’) |
|---|---|---|---|
| *Mx1-Cre* | CGTACTGACGGTGGGA GAAT | TGCATGATCTCCGGTATTGA | - |
| *Kat2a* | CACAGAGCTTCTTGGA GACC | GGCTTGATTCCTGTACCTCC | - |
| *Idh1* | GTTGGTGGATTCCATTGCTT | TGTTAGTCCCAACCCCTTCC | GACAAACTGACAGGCTG CAA |

| | | | |
|---|---|---|---|
| Amplification of U cassette | AAGTCCAACCTTATTGTCCCATCATAAGTTTTATACTCTGTAAGTAATGACCGCCTACTGCGACTATAGA | AGGTTCACCCTATGACTAACTGGCTCTAACAAAAGAGTTCTCAGCTCTTTAAGGCGCATAACGATACCAC | - |
| Amplification of G cassette | GCAATAGGAACCCTTTGCCATACTTAATTTTACTTCCATAAATCTCAAGTTCCTGTGTGAAATTGTTATCCGC | ACAAACTAGCTAACCTGATGGATGCAGTAATGAGTAACACAGGAGATCCTCCACTGGCCGTCGTTTTACA | - |
| Wildtype control assay | GGGCTAGGGGAAGCGCCATC | TGCGCAGGCCAAAGCCCAT | - |
| 5’ integration | TGGCTGGAAAACAAAAAGATCGG | CGTTATGCGCCTTAAAGAGCTGA | - |
| 3’ integration | TGGATCCGGGAAGTTCCTATTCC | TGGCACAGGCACAGAGGGA | - |
| 3’ internal probe | GGAGTGTTGTATCGCAGCAA | GCGCTAGGATTAAAGGCACA | - |
| 5’ internal probe | TCAGCATTCCCTAGGCACAA | TCTCTTGAGTGTGAGGCCAG | - |

**Pre-leukemia cell engraftment**

BM cells were isolated from long bones as described [12]. Briefly, following red blood cell lysis, BM nucleated cells were depleted of differentiated cells using a cocktail of biotinylated lineage (Lin) antibodies and streptavidin-labelled magnetic nanobeads (Biolegend), according to manufacturer’s instructions. Lin-depleted cells were cultured overnight at 37°C 5% $CO_2$ in RPMI supplemented with 20% Hi-FBS (R20), 2 mg/mL L-Glutamine, 1% PSA, 10 ng/mL of murine Interleukin 3 (mIL3), 10 ng/mL of murine Interleukin 6 (mIL6), and 20 ng/mL of murine Stem Cell Factor (mSCF) (cytokines from Peprotech) (supplemented R20), followed by retroviral transduction.

Retroviral construct *MSCV-AML1/ETO-IRES-GFP* was previously described [25]. Viral particle production and transduction were done as described previously [12]. GFP levels were assessed by flow cytometry. The BM cells obtained post transduction from *Kat2aWT* and *Kat2aNULL* animals were pooled individually, and one million cells were injected into >8 weeks old, C57/BL6 mice, which were lethally irradiated (2*5.5Gy). 17 mice/group were injected for leukemia studies. Leukemic mice were collected based on symptoms of hunched posture, inappetence and lethargy.

BM cells obtained from *Idh1*$^{R132H}$ transformed *Kat2aHET* and *Kat2aNULL* animals 20 weeks post pIpC were injected into CD45.1, C57/BL6 mice (n=8/group), which were sub-lethally irradiated (1*8Gy). Bones and spleens were collected post one year of transplantation and analyzed by flow cytometry.

**Colony forming cell assays**

For analysis of pre-leukemia samples from *RT1(9a)* and *Idh1*$^{R132H}$, 50,000 BM cells were plated in MethoCult M3434 (Stem Cell Technologies), following manufacturer's protocols. Colonies were scored 7-10 days after plating. Cells were collected from plates, washed and dispersed to a single-cell suspension, and serially re-plated for transformation analysis.

In S6K1 inhibition studies, 10,000 *Kat2a* WT BM cells transduced with *RT1(9a)* or carrying the recombined *Idh1*$^{R132H}$ allele, were plated in MethoCult M3434 containing freshly-added DMSO (vehicle) or 10µM PF4708671 (Tocris) with a final concentration of 0.1% DMSO. Colonies were scored as above.

**Leukemia maintenance *in vitro***

Pooled BM cells collected from 2-3 12-weeks old *Kat2a* floxed *Mx1-Cre* $^{-/-}$ animals without pIpC treatment, were retrovirally transduced with *RT1(9a)* and serially re-plated in MethoCult M3434 for a total of 3 platings (4000 cells/plating). At plate 3, cells were collected, transduced with a MIGR-Cre-OP-Puro (Cre+) or a MIGR-OP-Puro (empty) retrovirus, and cultured for 48 hours under puromycin selection, as described [25]. Transduced and antibiotic-selected cells were assessed for colony formation over two rounds of plating in Methocult M3434 (4000 cells/plating/condition) in the presence of puromycin, Colonies were scored 7-10 days after plating.

**Flow cytometry analysis**

Cell surface analysis of mouse BM and spleen was performed as described [12], using the antibodies in Table S2, with the following gating strategies: HSC- Lin$^{-}$ cKit$^{+}$ Sca1$^{+}$ CD34$^{-}$ Flt3$^{-}$; MPP-Lin$^{-}$ cKit$^{+}$ Sca1$^{+}$ CD34$^{+}$ Flt3$^{-}$; LMPP- Lin$^{-}$ cKit$^{+}$ Sca1$^{+}$ CD34$^{+}$ Flt3$^{+}$; CMP-Lin$^{-}$ cKit$^{+}$ Sca1$^{-}$ CD34$^{+/low}$ CD16/32$^{low}$; GMP- Lin$^{-}$ cKit$^{+}$ Sca1$^{+}$ CD34$^{+}$ CD16/32$^{high}$; MEP-Lin$^{-}$ cKit$^{+}$ Sca1$^{+}$ CD34$^{-}$ CD16/32$^{-}$; Lin$^{-}$- CD3e$^{-}$ B220$^{-}$ Gr1$^{-}$ CD11b$^{-}$ Ter119$^{-}$.

**Table S2: Antibodies used in flow cytometry analysis, cell sorting and lineage selection**

| Antibody | Fluorochrome | Catalogue ID | Clone | Dilution | Supplier |
|---|---|---|---|---|---|
| CD45R/ B220 | APC-Cy7 | 103223 | RA3-6B2 | 1:50 | BioLegend |
| CD45R/ B220 | PerCP-Cy5.5 | 103235 | RA3-6B2 | 1:100 | BioLegend |
| CD117/c-Kit | APC-Cy7 | 105826 | 2B8 | 1:50 | BioLegend |
| CD11b/Mac1 | AF700 | 101222 | M1/70 | 1:200 | BioLegend |
| CD16/32/FcγR | PE | 101308 | 93 | 1:100 | BioLegend |
| CD34 | APC | 128612 | HM34 | 1:100 | BioLegend |
| F4/80 | PE | 123109 | BM8 | 1:100 | BioLegend |
| Gr1 | PB | 108430 | RB6-8C5 | 1:100 | BioLegend |
| Hoechst 33258 | - | H3569 | - | 1:10000 | Invitrogen |
| Sca1 | PE-Cy7 | 108114 | D7 | 1:100 | BioLegend |
| Streptavidin | BV421 | 405226 | - | 1:200 | BioLegend |
| Streptavidin | BV605 | 405229 | - | 1:200 | BioLegend |
| CD45R/B220 (Lin) | Biotin | 103204 | RA3-6B2 | 1:300 | BioLegend |
| Ter119 (Lin) | Biotin | 116204 | Ter119 | 1:300 | BioLegend |
| Gr1 (Lin) | Biotin | 108404 | RB6-8C5 | 1:300 | Biolegend |
| CD3e (Lin) | Biotin | 100304 | 145–2 C11 | 1:300 | BioLegend |
| CD11b (Lin) | Biotin | 101204 | M1/70 | 1:300 | BioLegend |
| Nanobeads | Streptavidin | 76447 | - | 1:10 | BioLegend |
| Click-iT Cell Reaction Buffer Kit | AF647 azide | A10277 | - | 1:500 | Invitrogen |

**Single-cell RNA sequencing and analysis**

Pre-leukemia BM samples were collected from individual animals engrafted with *RT1(9a)*-transduced *Kat2aWT* and *Kat2aNULL* cells, 2 and 4 months after transplantation, and stored at -150°C. Cells were thawed, recovered in R20 medium, and sorted on an Influx sorter (BD) as Hoechst 33258-negative (live), $GFP^+$ (RT1*(9a)* reporter), $cKit^+$ (early progenitors) singlets. Sorted cells were immediately used for library preparation with Chromium Next GEM Single Cell 3’GEM, Library and Gel Bead Kit v2 (10XGenomics). Libraries were quality-controlled and underwent paired-end sequencing on an Illumina NextSeq 500 Sequencer. Library preparation and sequencing were performed at CRUK Cambridge Research Institute. Raw single cell RNAseq fastq reads were analysed using Cellranger software (v2.2) to obtain the cell-gene count-matrix (Table S3). The count-matrix data was pre-processed with Seurat v2.4 [26] as described [12]. Differential gene expression was obtained with DESeq2 [27]. Gene ontology analysis was performed using Panther 14.0 [28] selecting Fischer's exact test with Bonferroni correction. Transcriptional variability analysis used pairwise distance between gene correlations as a measure of cellular heterogeneity, by identifying the top 500 highly variable genes based on distance-to-median (DM) and calculating Spearman correlation coefficients between all gene pairs [19]. Pseudotime analysis was performed using Monocle v3.0 [21] individually for *Kat2aWT* and *Kat2aNULL* cells.

**Table S3: Cell-Gene count matrix specification**

| | |
|---|---|
| Total number of cells sequenced | 1767 |
| Median number of genes per cell | 1575 |
| *Kat2aWT* 2 months | 379 |
| *Kat2aNULL* 2 months | 369 |
| *Kat2aWT* 4 months | 518 |
| *Kat2aNULL* 4 months | 501 |

**Bulk RNA-sequencing analysis**

Total RNA was extracted from mouse bone marrow aspirates, following enucleated cell lysis. Paired-end RNA-seq reads were mapped to the mouse genome (UCSC mm10) using STAR with default parameters [29]. The total number of reads aligning to the exons of each gene (as per GENCODE vM4, [30]) were counted using HTSeq-count [31]. Read counts were used for FPKM computation and differential expression analyses using DESeq2 between sex matched pre-leukemia and AML samples [27]. Pre-leukemia samples; *Idh1*$^{R132H}$ (n=2), *Idh1*$^{R132H}$ *Npm1c* (n=2), *Npm1c N-Ras*$^{G12D}$ *Idh1*$^{R132H}$ (n=2), *Idh1*$^{WT}$(n=6 female, 4 male) and AML samples; *N-Ras*$^{G12D}$ *Idh1*$^{R132H}$ (n=3), *Npm1c N-Ras*$^{G12D}$ *Idh1*$^{R132H}$ (n=3).

**Measurement of protein synthesis**

Protein synthesis rates were estimated using O-propargyl-puromycin (OP-Puro, Thermo Fisher Scientific) incorporation, as described [12]. In detail, one million *RT1(9a) Kat2aWT* or *Idh1*$^{R132H}$

*Kat2aHET* cells treated with S6K1 inhibitor PF4708671 10µM vs. DMSO (vehicle) were collected from successive re-plating of colony-forming assays and cultured for 2 hours in the presence of cytokines, mSCF (20ng/ml), mIL-3 (10ng/ml) and mIL-6 (10ng/ml) in R20 medium. In other assays, $Idh1^{R132H}$ *Kat2aHET* vs. *Kat2aNULL* BM cells collected 20 weeks after pIpC treatment for locus activation / excision, were thawed and cultured overnight in the same conditions. A final concentration of 12.5 µM OP-Puro was added directly to 80% of each culture for the last hour of the culture period; the remainder 20% cells were treated with PBS and processed in parallel as control. After incubation, cells were washed with ice cold PBS without $Ca^{2+}$ or $Mg^{2+}$ (Sigma), and resuspended in PBS/10%FBS for cell surface staining with c-Kit-APCC7, CD11b-Biotin, Gr1-Biotin (all from BioLegend, see Table 2), followed by Streptavidin Brilliant Violet 605 (also from BioLegend) both staining steps for 30 min on ice. After washing, cells were fixed in 1% paraformaldehyde (PFA) in PBS for 15 min on ice protected from light, washed, and permeabilized in PBS/3% FBS/0.1% saponin (permeabilization buffer) at room temperature, in the dark, for 5 min. Cells were washed and used immediately in the azide-alkyne cyclo-addition reaction with Click-iT Cell Reaction Buffer Kit (Thermo Fisher Scientific; C10269) and Alexa Fluor 647-Azide (Thermo Fisher Scientific; A10277) with a master reaction solution freshly prepared for immediate use, as per manufacturer's instructions. Alexa Fluor 647-Azide was used at a final concentration of 5µM. The reaction proceeded in the dark at room temperature for 30 min; cells were washed twice in permeabilization buffer and resuspended in PBS, 5 minutes prior to flow cytometry analysis.

**Statistical analysis**

Experiments were performed in triplicate, with any exceptions specifically indicated in the text or Figure Legends. Data are plotted as Mean ± Standard deviation with statistical tests described in the respective Figure Legends. Statistical analysis was performed using Graphpad Prism 8.0 software. R language was used for analysis of single-cell RNA-seq data.

### SUPPLEMENTARY FIGURES

Fig S1

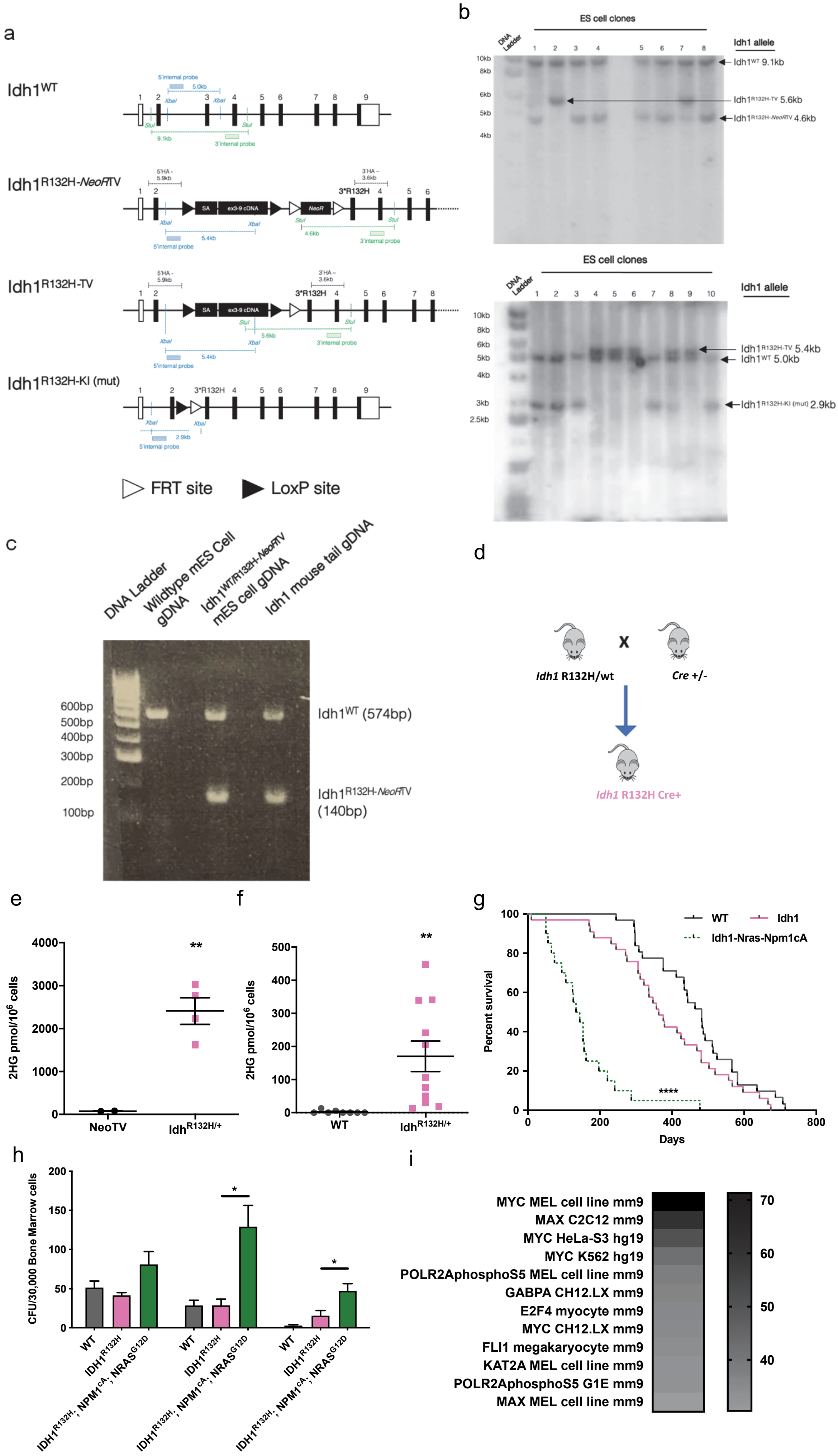
a
Idh1WT
Idh1R132H-NeoRTV
Idh1R132H-TV
Idh1R132H-KI (mut)
FRT site
LoxP site
b
ES cell clones
Idh1 allele
Idh1WT 9.1kb
Idh1R132H-TV 5.6kb
Idh1R132H-NeoRTV 4.6kb
Idh1R132H-TV 5.4kb
Idh1WT 5.0kb
Idh1R132H-KI (mut) 2.9kb
c
DNA Ladder
Wildtype mES Cell gDNA
Idh1WT/R132H-NeoRTV mES cell gDNA
Idh1 mouse tail gDNA
600bp
500bp
400bp
300bp
200bp
100bp
Idh1WT (574bp)
Idh1R132H-NeoRTV (140bp)
d
Idh1 R132H/wt
x
Cre +/-
Idh1 R132H Cre+
e
2HG pmol/10⁶ cells
NeoTV
IdhR132H/+
**
f
2HG pmol/10⁶ cells
WT
IdhR132H/+
**
g
Percent survival
Days
WT
Idh1
Idh1-Nras-Npm1cA
****
h
CFU/30,000 Bone Marrow cells
WT
IDH1R132H
IDH1R132H; NPM1cA; NRASG12D
*
i
MYC MEL cell line mm9
MAX C2C12 mm9
MYC HeLa-S3 hg19
MYC K562 hg19
POLR2AphosphoS5 MEL cell line mm9
GABPA CH12.LX mm9
E2F4 myocyte mm9
MYC CH12.LX mm9
FLI1 megakaryocyte mm9
KAT2A MEL cell line mm9
POLR2AphosphoS5 G1E mm9
MAX MEL cell line mm9

**Fig. S1: Development and analysis of the *Idh1*$^{R132H}$ pre-leukemia model. (A)** Schematic of *Idh1*$^{WT}$ and stepwise assembly of *Idh1*$^{R132H}$ knock-in (KI) mutant (mut) allele, with representation of Southern Blotting probes used for confirmation of successful generation of intermediate *Idh1*$^{R132H-NeoR-TV}$ and *Idh1*$^{R132H-TV}$ alleles, as represented in panel B. **(B** Southern Blots confirming excision of the NeoR resistance cassette and LoxP recombination to generate *Idh1*$^{R132H-NeoR-TV}$ and *Idh1*$^{R132H-TV}$ alleles. **(C)** Genotyping agarose gel for selection of F1 mice carrying an inducible *Idh1*$^{R132H-NeoR-TV}$ allele after backcross to *RosaFLPe*-negative mice, as described in Supplementary Methods. **(D)** Schematic of*Idh1*$^{R132H}$ and *Mx1-Cre* mouse crosses to generate a *Cre* inducible *Idh1*$^{R132H}$ mouse. **(E)** Mass spectrometry (MS) quantification of 2-Hydroxyglutarate (2-HG) in control (NeoTV) and *Idh1*$^{R132H}$ (*Idh*$^{R132H/+}$) embryonic stem cells; mean ± SD, n=2-4 samples/condition. **(F)** MS quantification of 2-HG in control (WT) and *Idh1*$^{R132H}$ (*Idh*$^{R132H/+}$*)* bone marrow (BM) cells; mean ± SD, n=8-11 animals/genotype, **p<0.01, 2-tailed t-test. **(G)** Survival curve of mice with *Idh1*$^{R132H}$ or *Idh1*$^{R132H}$*-Nras*$^{G12D}$*-Npm1cA* alleles; n= 20-33 animals/genotype, ****p<0.0001, log-rank test. **(H)** Colony-forming assay of BM cells obtained from WT, *Idh1*$^{R132H}$*, Idh1R132H-Npm1*$^{cA}$*- Nras*$^{G12D}$ animals, with serial re-plating; mean ± SD, n=4-10/genotype, *p<0.05, 2-tailed t-test. **(I)** EnrichR [32] analysis of Encode transcription factor binding enrichment in differentially down-regulated genes in *Idh1*$^{R132H}$ leukemia vs pre-leukemia samples; *p-adj<0.05. RNA-seq analysis of *Idh1*$^{R132H}$ and *Idh1R132H-Npm1*$^{cA}$*-NRAS*$^{G12D}$ pre-leukemia (Lin$^{-}$ cells collected 4 weeks after pIpC treatment) vs. *Idh1R132H-Npm1*$^{cA}$*-NRAS*$^{G12D}$ leukemia samples (terminal).

Fig S2

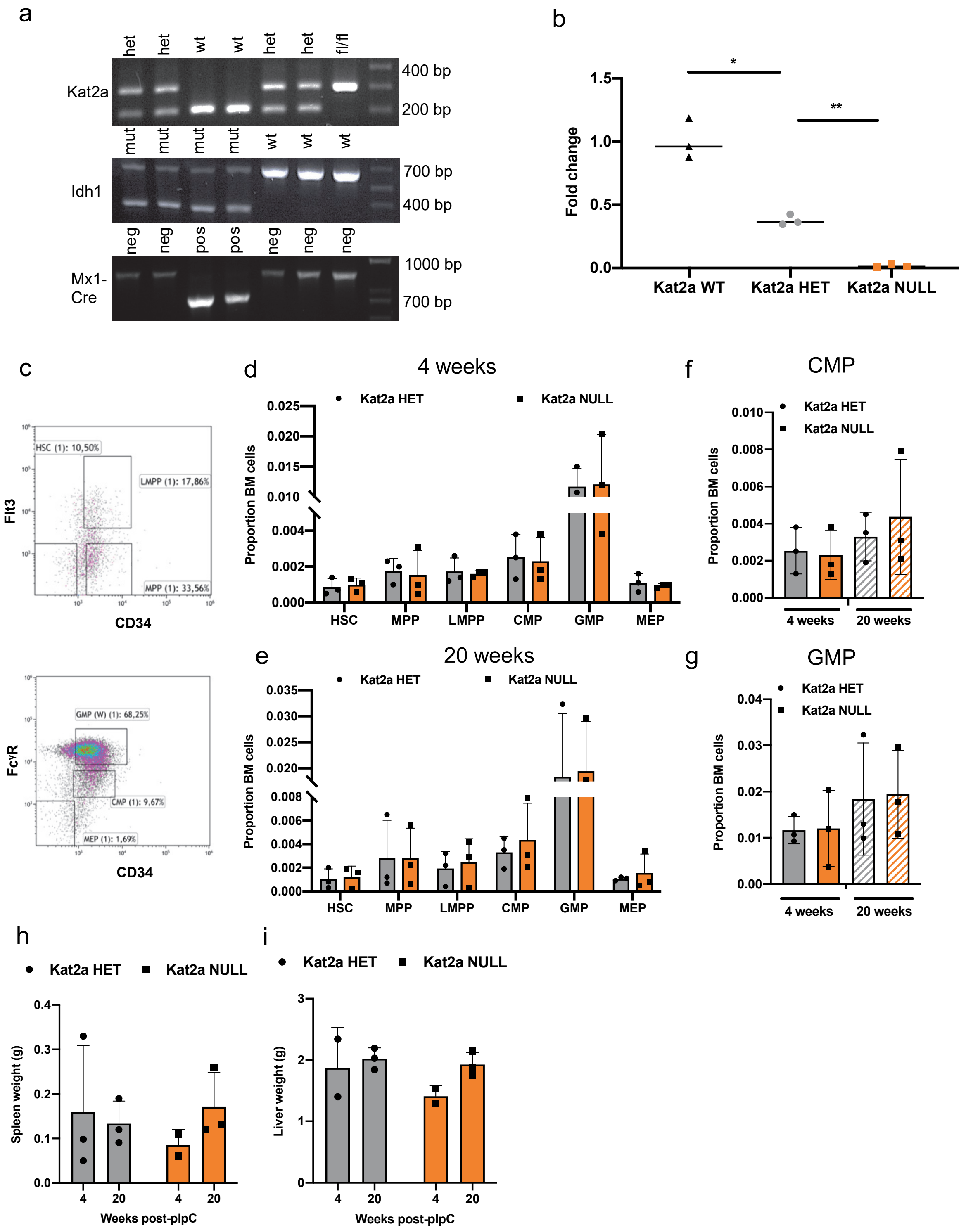
a
het
het
wt
wt
het
het
fl/fl
Kat2a
400 bp
200 bp
mut
mut
mut
mut
wt
wt
wt
Idh1
700 bp
400 bp
neg
neg
pos
pos
neg
neg
neg
Mx1-Cre
1000 bp
700 bp
b
Fold change
Kat2a WT
Kat2a HET
Kat2a NULL
*
**
c
HSC (1): 10,50%
LMPP (1): 17,86%
MPP (1): 33,56%
Flt3
CD34
GMP (W) (1): 68,25%
CMP (1): 9,67%
MEP (1): 1,69%
FcγR
CD34
d
4 weeks
Kat2a HET
Kat2a NULL
Proportion BM cells
HSC
MPP
LMPP
CMP
GMP
MEP
e
20 weeks
Kat2a HET
Kat2a NULL
Proportion BM cells
HSC
MPP
LMPP
CMP
GMP
MEP
f
CMP
Kat2a HET
Kat2a NULL
Proportion BM cells
4 weeks
20 weeks
g
GMP
Kat2a HET
Kat2a NULL
Proportion BM cells
4 weeks
20 weeks
h
Kat2a HET
Kat2a NULL
Spleen weight (g)
Weeks post-pIpC
i
Kat2a HET
Kat2a NULL
Liver weight (g)
Weeks post-pIpC

**Fig. S2: Cellular characterization of *Idh1*$^{R132H}$ pre-leukemia. (A)** Representative PCR genotyping gel images of *Kat2a*, *Idh1* and *Mx1-Cre* alleles. **(B)** qRT-PCR analysis of *Kat2a* expression in BM cells of *Kat2aWT*, *Kat2aHET* and *Kat2aNULL* animals; mean ± SD, n=3 animals/genotype, *p<0.05, **p<0.01, 2-tailed t-test. **(C)** Flow cytometry gating strategy for enumeration of BM stem and progenitor cells in *Idh1* mut *Kat2a* HET and *Idh1* mut *Kat2a* NULL pre-leukemic animals. **(D)** Proportion of BM stem and progenitor cells in *Idh1* mut *Kat2a* HET and *Idh1* mut *Kat2a* NULL pre-leukemic animals 4 weeks post-pIpC treatment; mean ± SD, n=3, 2-tailed t-test. **(E)** Proportion of BM stem and progenitor cells in *Idh1* mut *Kat2a* HET and *Idh1* mut *Kat2a* NULL pre-leukemic animals 20-week post-pIpC treatment; mean ± SD, n=3, 2-tailed t-test. **(F)** Proportion of BM cells characterized as CMP population of cells at 4 week and 20 week time point; mean ± SD, n=3, 2-tailed t-test. **(G)** Proportion of BM cells characterized as GMP population of cells at 4 week and 20 week time point; mean ± SD, n=3, 2-tailed t-test. (**H-I)** Analysis of pre-leukemia burden 4 and 20-weeks post-pIpC, **(H)** spleen weights, **(I)** liver weights; mean ± SD, n=2-3.

Fig S3

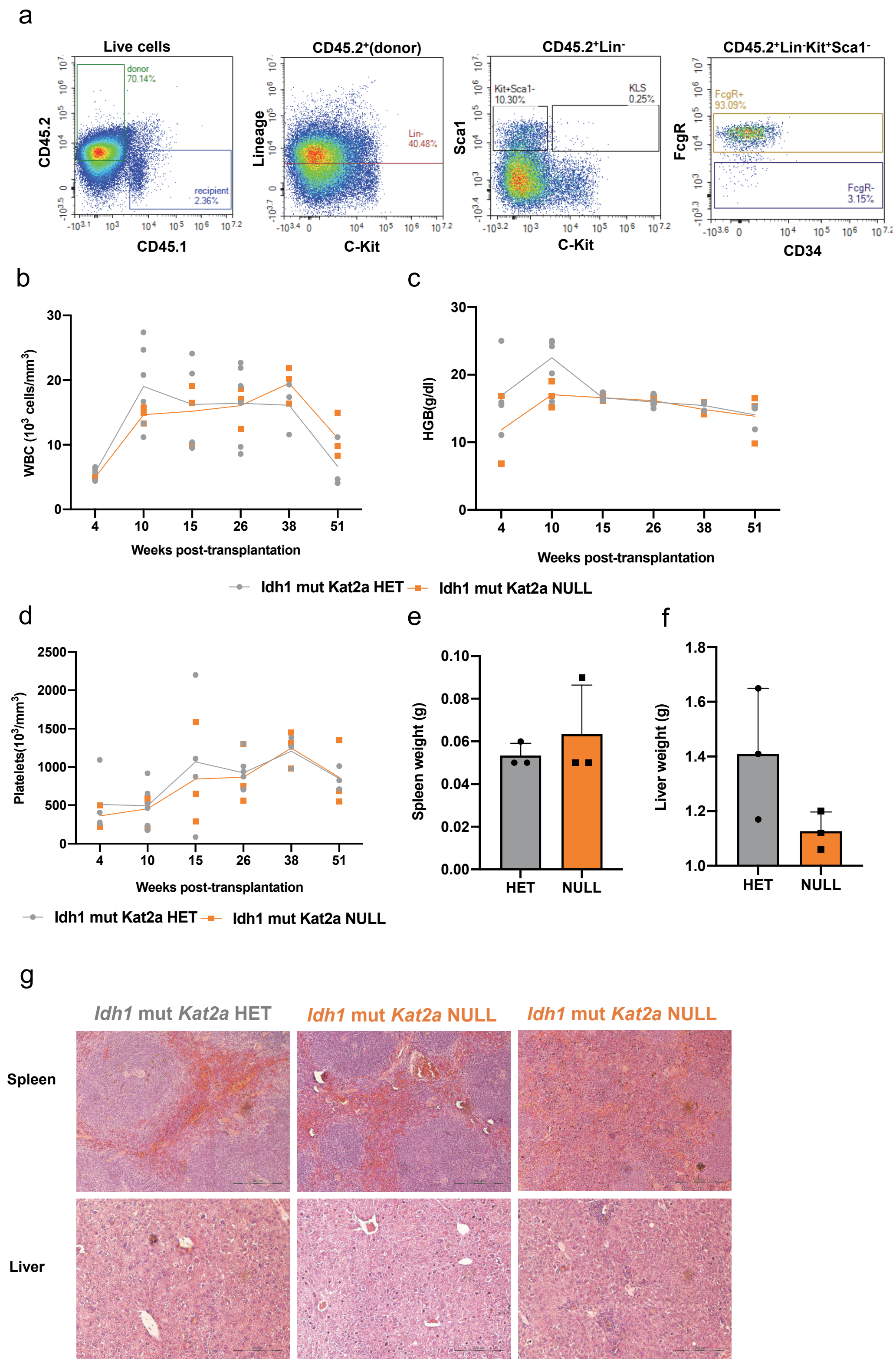
a
Live cells
CD45.2
CD45.1
donor 70.14%
recipient 2.36%
CD45.2+(donor)
Lineage
C-Kit
Lin- 40.48%
CD45.2+Lin-
Sca1
C-Kit
Kit+Sca1- 10.30%
KLS 0.25%
CD45.2+Lin-Kit+Sca1-
FcgR
CD34
FcgR+ 93.09%
FcgR- 3.15%
b
WBC (10^3 cells/mm^3)
Weeks post-transplantation
c
HGB(g/dl)
Weeks post-transplantation
Idh1 mut Kat2a HET
Idh1 mut Kat2a NULL
d
Platelets(10^3/mm^3)
Weeks post-transplantation
Idh1 mut Kat2a HET
Idh1 mut Kat2a NULL
e
Spleen weight (g)
HET
NULL
f
Liver weight (g)
HET
NULL
g
Idh1 mut Kat2a HET
Idh1 mut Kat2a NULL
Idh1 mut Kat2a NULL
Spleen
Liver

**Fig. S3: Investigation of *Idh1*$^{R132H}$ pre-leukemia transplants. (A)** Flow cytometry gating strategy for engrafted *Idh1* mut *Kat2a* HET and *Idh1* mut *Kat2a* NULL BM cells. **(B-D)** Analysis of hematological parameters from peripheral blood of mice engrafted with *Idh1* mut *Kat2a* HET and *Idh1* mut *Kat2a* NULL BM cells: **(B)** WBC, **(C)** Hemoglobin, **(D)** Platelets; mean ± SD, n=3-6 at 10 week time point, n.s. No animals (6 receiving *Idh1* mut *Kat2a* HET / 3 receiving *Idh1* mut *Kat2a* NULL cells) developed signs or symptoms or leukemia during the observation period. Three animals were culled for welfare reasons unrelated to a leukemia disease process. **(E-F)** Analysis of leukemia burden 1-year post-transplantation: **(E)** spleen weight, **(F)** liver weight; mean ± SD, n=3/genotype, 2-tailed t-test. **(G)** Histology analysis of *Idh1* mut *Kat2a* HET and *Idh1* mut *Kat2a* NULL recipient animals culled 1-year post-transplantation.

Fig S4

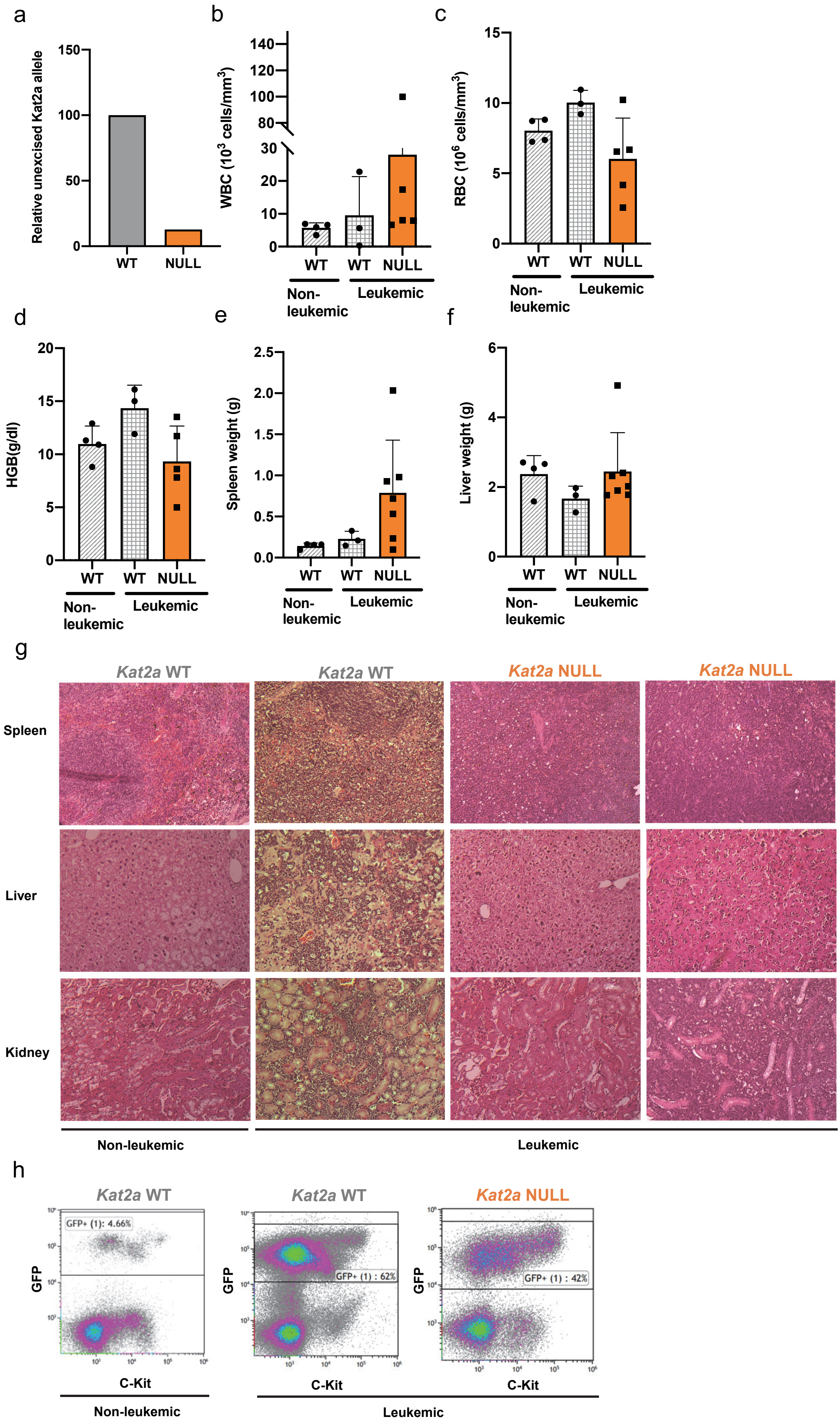

a
Relative unexcised Kat2a allele
150
100
50
0
WT
NULL
b
WBC (10^3 cells/mm^3)
140
120
100
80
30
20
10
0
WT
WT
NULL
Non-leukemic
Leukemic
c
RBC (10^6 cells/mm^3)
15
10
5
0
WT
WT
NULL
Non-leukemic
Leukemic
d
HGB(g/dl)
20
15
10
5
0
WT
WT
NULL
Non-leukemic
Leukemic
e
Spleen weight (g)
2.5
2.0
1.5
1.0
0.5
0.0
WT
WT
NULL
Non-leukemic
Leukemic
f
Liver weight (g)
6
4
2
0
WT
WT
NULL
Non-leukemic
Leukemic
g
Kat2a WT
Kat2a WT
Kat2a NULL
Kat2a NULL
Spleen
Liver
Kidney
Non-leukemic
Leukemic
h
Kat2a WT
Kat2a WT
Kat2a NULL
GFP+ (1): 4.66%
GFP+ (1) : 62%
GFP+ (1) : 42%
GFP
C-Kit
Non-leukemic
Leukemic

**Fig. S4: Characterization of *RT1(9a) Kat2a* NULL leukemia. (A)** Relative expression of *Kat2a* in *RT1(9a)*-transformed *Kat2aWT* and *Kat2aNULL* BM cells; transformation used pools of 3 animals/genotype. **(B-D)** Terminal peripheral blood analysis of mice engrafted with *RT1(9a)*-transduced *Kat2aWT* and *Kat2aNULL* $Kit^+$ / $Lin^-$ cells and followed up for leukemia development. Animals were analysed upon exhibition of terminal symptoms, or after 405 days, if asymptomatic. **(B)** WBC, **(C)** RBC, **(D)** HGB; mean ± SD, n=3-5, 2-tailed t-test. **(E-F)** Analysis of leukemia burden at terminal time point in the same animals. **(E)** Spleen weight; **(F)** liver weight. **(G)** Representative histology images of spleen, liver, and kidney tissues of the same engrafted mouse cohort at terminal time point, with or without leukemia development. **(H)** Representative flow cytometry plots of terminal *Kat2aWT* or *Kat2aNULL RT1(9a)*-engrafted mice, with or without leukemia development.

Fig S5

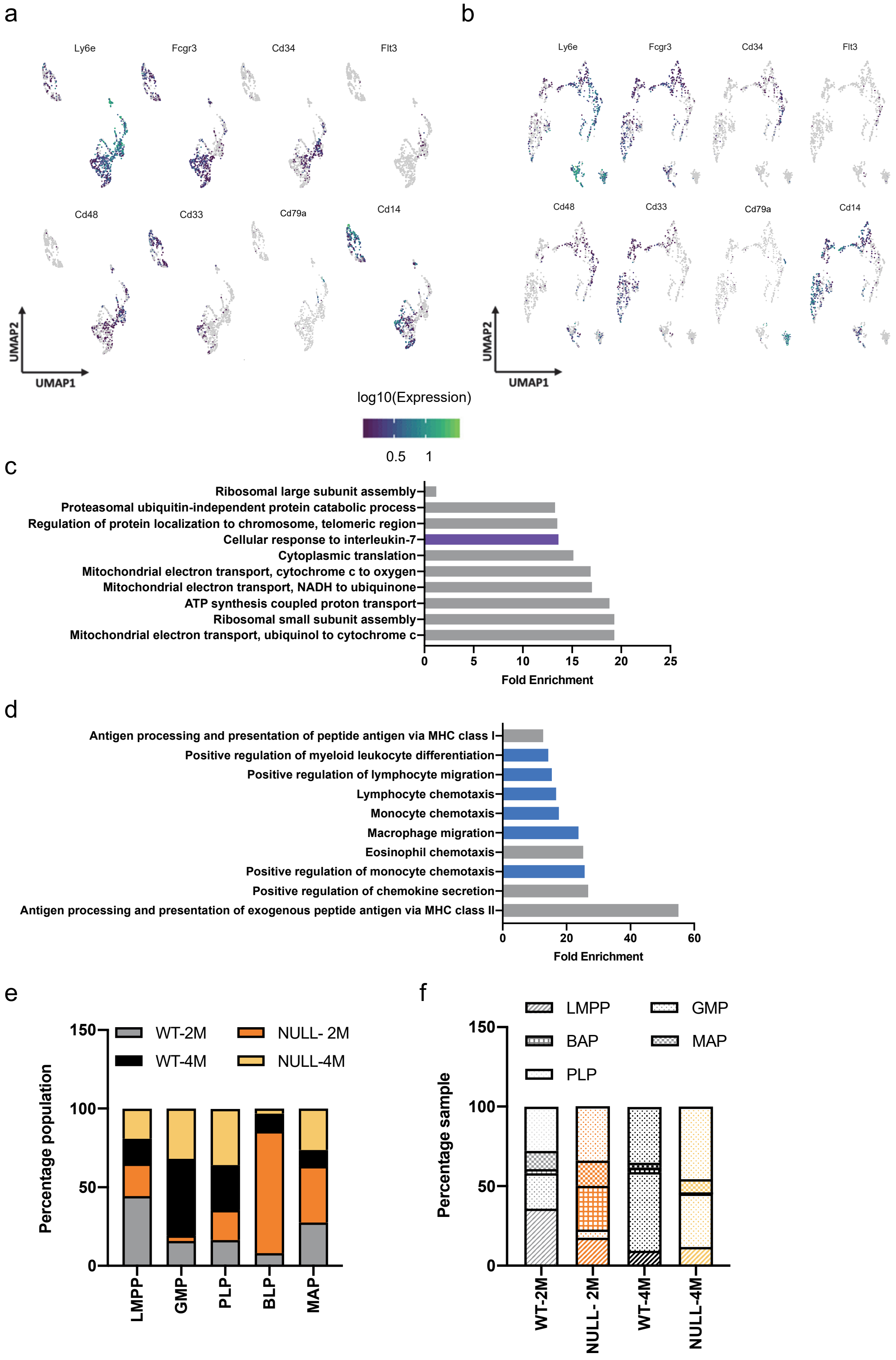
a
Ly6e
Fcgr3
Cd34
Flt3
Cd48
Cd33
Cd79a
Cd14
UMAP2
UMAP1
b
Ly6e
Fcgr3
Cd34
Flt3
Cd48
Cd33
Cd79a
Cd14
UMAP2
UMAP1
log10(Expression)
0.5
1
c
Ribosomal large subunit assembly
Proteasomal ubiquitin-independent protein catabolic process
Regulation of protein localization to chromosome, telomeric region
Cellular response to interleukin-7
Cytoplasmic translation
Mitochondrial electron transport, cytochrome c to oxygen
Mitochondrial electron transport, NADH to ubiquinone
ATP synthesis coupled proton transport
Ribosomal small subunit assembly
Mitochondrial electron transport, ubiquinol to cytochrome c
0
5
10
15
20
25
Fold Enrichment
d
Antigen processing and presentation of peptide antigen via MHC class I
Positive regulation of myeloid leukocyte differentiation
Positive regulation of lymphocyte migration
Lymphocyte chemotaxis
Monocyte chemotaxis
Macrophage migration
Eosinophil chemotaxis
Positive regulation of monocyte chemotaxis
Positive regulation of chemokine secretion
Antigen processing and presentation of exogenous peptide antigen via MHC class II
0
20
40
60
Fold Enrichment
e
WT-2M
NULL- 2M
WT-4M
NULL-4M
Percentage population
150
100
50
0
LMPP
GMP
PLP
BLP
MAP
f
LMPP
GMP
BAP
MAP
PLP
Percentage sample
150
100
50
0
WT-2M
NULL- 2M
WT-4M
NULL-4M

**Fig. S5: Representation of hematopoietic lineage-associated surface markers and regulatory transcription factors on RT1(9a) pseudotime trajectories. (A-B)** Expression of lineage-affiliated hematopoietic markers in single-cell trajectories of RT1(9a) transduced BM; **(A)** *Kat2aWT*, **(B)** *Kat2aNULL* cells. *Ly6e*: hematopoietic stem cell (HSC), multipotent (MPP) and lympho/myeloid (LMPP) progenitors; myeloid cells). *Fcgr3*; myeloid progenitors, myelo-monocytic cells. *Cd34*: LMPP, MPP, myeloid progenitors, multipotent and lympho/myeloid progenitor marker), *Flt3*; MPP, LMPP, lymphoid cells. *Cd48*: committed progenitors and lymphocytes. *Cd33*; myeloid cells. *Cd14*; monocyte/macrophage. *Cd79a*: B lymphocytes. **(C)** Gene ontology analysis of genes overexpressed in B-affiliated progenitors(BAP) using Panther14.0 [28]; *p-adj<0.05. **(D)** Gene ontology analysis of genes overexpressed in monocyte-affiliated progenitors (MAP) using Panther14.0; *p-adj<0.05. **(E)** Proportion of candidate progenitor cell compartments in the RT1(9a) pseudotime trajectory contributed by individual *Kat2aWT* or *Kat2aNULL*, 2-month or 4-month samples. LMPP: lymphoid/myeloid-primed progenitor; GMP: granulocyte-monocyte progenitor; MAP: monocyte-affiliated progenitor; BAP: B-cell affiliated progenitor; PLP: Pre-leukemia progenitor. **(F)** Proportion of RT1(9a)-transduced cells in each individual time-point sample (*Kat2aWT* or *Kat2a*NULL, 2 or 4 months post-engraftment) contributed by candidate progenitor compartments captured by pseudotime trajectory analysis.

Fig S6

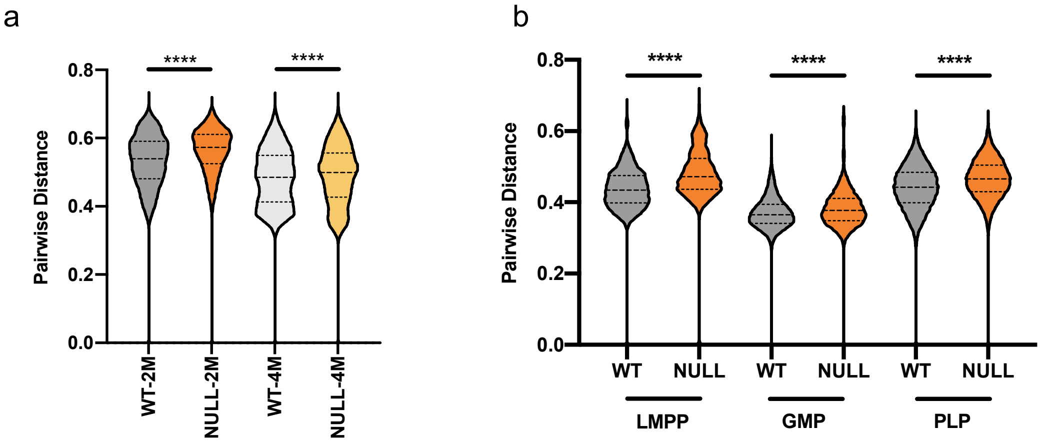
a
Pairwise Distance
0.8
0.6
0.4
0.2
0.0
****
****
WT-2M
NULL-2M
WT-4M
NULL-4M
b
Pairwise Distance
0.8
0.6
0.4
0.2
0.0
****
****
****
WT
NULL
WT
NULL
WT
NULL
LMPP
GMP
PLP

**Fig. S6: Loss of *Kat2a* and transition into pre-leukemia associate with enhanced cell-to-cell transcriptional variability of RT1(9a) cells. (A)** Comparison of pairwise distances between individual *Kat2aWT* and *Kat2aNULL* RT1(9a) cells measured at 2 and 4 months post-engraftment. Pairwise distance measure as in Fig. 3G. ****p-adj<0.0001, 2-tailed t-test. **(B)** Comparison of RT1(9a) *Kat2aWT* and *Kat2aNULL* genotype-specific pairwise distances within individual LMPP, GMP, andPLP compartments; all ****p-adj<0.0001, 2-tailed t-test.

Fig S7

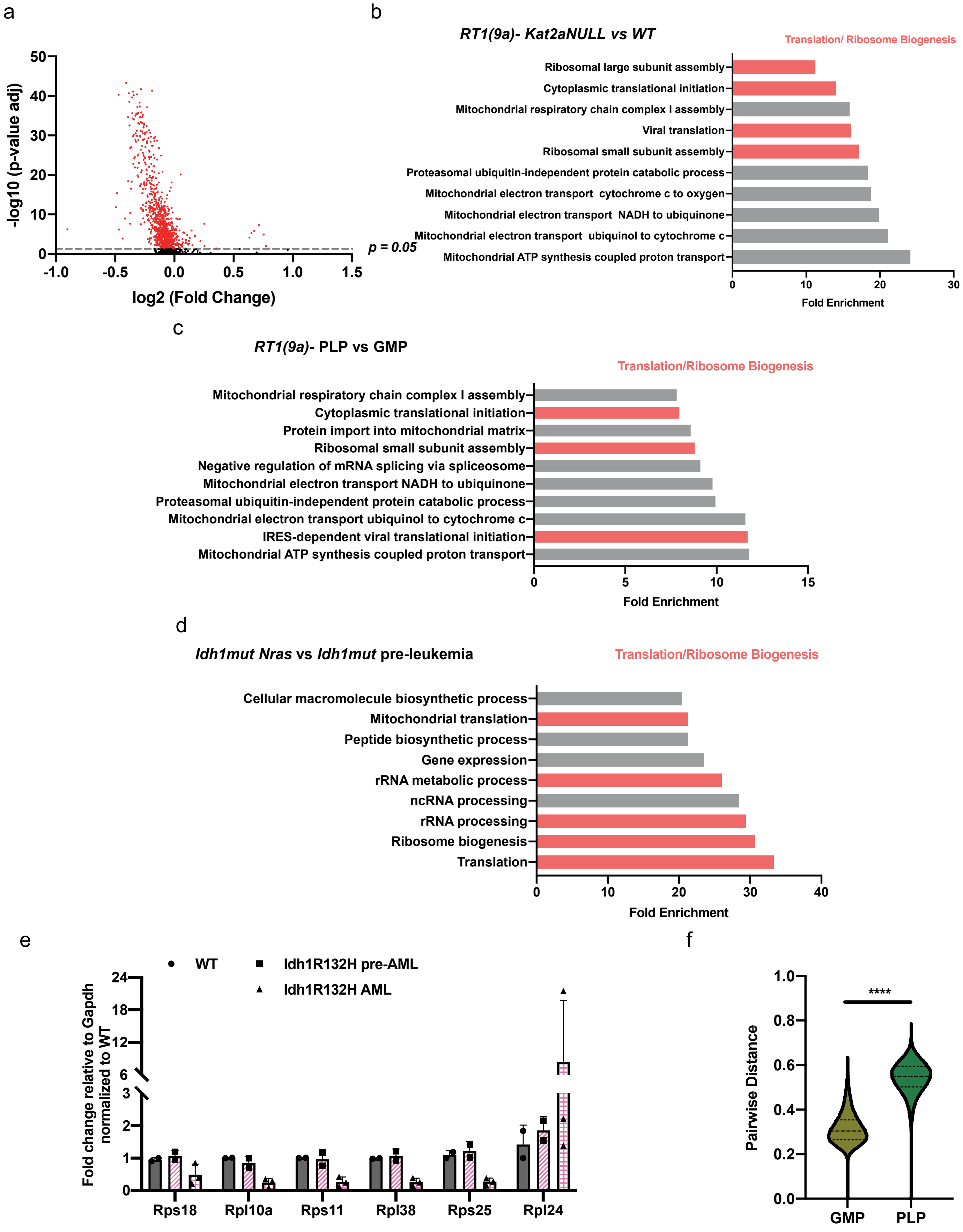
a
-log10 (p-value adj)
50
40
30
20
10
0
-1.0
-0.5
0.0
0.5
1.0
1.5
log2 (Fold Change)
p = 0.05
b
RT1(9a)- Kat2aNULL vs WT
Translation/ Ribosome Biogenesis
Ribosomal large subunit assembly
Cytoplasmic translational initiation
Mitochondrial respiratory chain complex I assembly
Viral translation
Ribosomal small subunit assembly
Proteasomal ubiquitin-independent protein catabolic process
Mitochondrial electron transport cytochrome c to oxygen
Mitochondrial electron transport NADH to ubiquinone
Mitochondrial electron transport ubiquinol to cytochrome c
Mitochondrial ATP synthesis coupled proton transport
0
10
20
30
Fold Enrichment
c
RT1(9a)- PLP vs GMP
Translation/Ribosome Biogenesis
Mitochondrial respiratory chain complex I assembly
Cytoplasmic translational initiation
Protein import into mitochondrial matrix
Ribosomal small subunit assembly
Negative regulation of mRNA splicing via spliceosome
Mitochondrial electron transport NADH to ubiquinone
Proteasomal ubiquitin-independent protein catabolic process
Mitochondrial electron transport ubiquinol to cytochrome c
IRES-dependent viral translational initiation
Mitochondrial ATP synthesis coupled proton transport
0
5
10
15
Fold Enrichment
d
Idh1mut Nras vs Idh1mut pre-leukemia
Translation/Ribosome Biogenesis
Cellular macromolecule biosynthetic process
Mitochondrial translation
Peptide biosynthetic process
Gene expression
rRNA metabolic process
ncRNA processing
rRNA processing
Ribosome biogenesis
Translation
0
10
20
30
40
Fold Enrichment
e
WT
Idh1R132H pre-AML
Idh1R132H AML
Fold change relative to Gapdh normalized to WT
24
18
12
6
3
2
1
0
Rps18
Rpl10a
Rps11
Rpl38
Rps25
Rpl24
f
Pairwise Distance
1.0
0.8
0.6
0.4
0.2
0.0
****
GMP
PLP

**Fig. S7: *Kat2a* loss downregulates protein synthesis genes in pre-leukemia progression. (A)** Volcano plot of single-cell RNA-seq data for RT1(9a) *Kat2aNULL* vs *WT* cells. Differentially expressed genes at p-adj <0.05 shown in red. **(B)** Over-represented gene ontologies (GO) for genes downregulated in RT1(9a) *Kat2aNULL* vs *WT* cells. Panther14.0 analysis of single-cell RNA-seq data, with 2 and 4-month timepoints analysed together; *p-adj<0.05. **(C)** Over-represented GO for genes down-regulated in RT1(9a) *Kat2aWT* PLP vs GMP *p-adj<0.05. **(D)** Over-represented GO for genes down-regulated in *N-Ras*$^{G12D}$ *Idh1*$^{R132H}$ and *Npm1c N-Ras*$^{G12D}$ *Idh1*$^{R132H}$ AML compared to *Idh1*$^{R132H}$, *Idh1*$^{R132H}$ *Npm1c*, and *Npm1c N-Ras*$^{G12D}$ *Idh1*$^{R132H}$ pre-leukemia; *p-adj<0.05. Genes were not differential in *Idh1*$^{R132H}$ vs *Idh1*$^{WT}$, or in *Idh1*$^{R132H}$ *Npm1c* and *Npm1c N-Ras*$^{G12D}$ *Idh1*$^{R132H}$ vs *Idh1*$^{R132H}$ pre-leukemia. **(E)** qRT-PCR analysis of ribosomal protein genes in *Idh1*$^{R132H}$ leukemia vs. pre-leukemia samples; mean ± SD, n=2-3. Data from Lin$^{-}$ BM plotted relative to *Idh1* WT Lin$^{-}$ cells, normalised for *Gapdh* expression. **(F)** Comparison of pairwise distances between individual cells in GMP and PLP compartments using the ribosomal biogenesis signature in Fig. 6D-E; ****p-adj<0.0001, 2-tailed t-test.

Fig S8

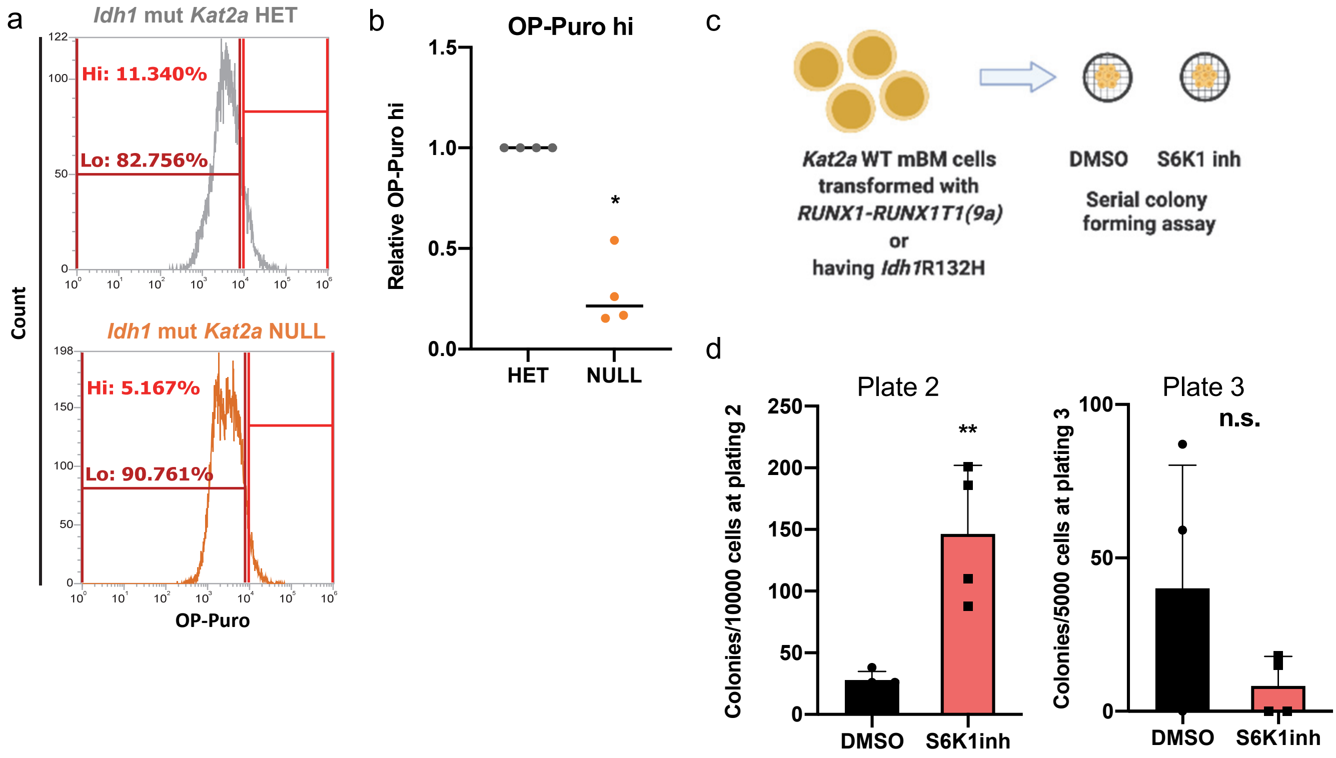
a
Idh1 mut Kat2a HET
Hi: 11.340%
Lo: 82.756%
Idh1 mut Kat2a NULL
Hi: 5.167%
Lo: 90.761%
Count
OP-Puro
b
OP-Puro hi
Relative OP-Puro hi
1.5
1.0
0.5
0.0
*
HET
NULL
c
Kat2a WT mBM cells
transformed with
RUNX1-RUNX1T1(9a)
or
having Idh1R132H
DMSO
S6K1 inh
Serial colony
forming assay
d
Plate 2
Colonies/10000 cells at plating 2
250
200
150
100
50
0
**
DMSO
S6K1inh
Plate 3
Colonies/5000 cells at plating 3
100
50
0
n.s.
DMSO
S6K1inh

**Fig. S8: Inhibition of protein synthesis facilitates pre-leukemia transformation.** (**A**) Representative flow cytometry plots of OP-Puro incorporation in *Kat2aHET* vs. *Kat2aNULL* $Idh1^{R132H}$ BM cells collected 20 weeks post-pIpC and treated *in vitro*. **(B)** Quantification of OP-Puro high $Idh1^{R132H}$ *Kat2aHET* and *Kat2aNULL* cells represented in A; mean ± SD, n=4, *p<0.05, 2-tailed t-test. (**C**) Schematic of S6K1 inhibition assays. **(D)** CFC replating of $Idh1^{R132H}$ *Kat2aWT* cells in the presence of S6K1inh (or DMSO), 4 weeks post-locus activation. Plate 2 (left); mean ± SD, n=4, **p<0.01, Plate 3 (right); mean ± SD, n=4, n.s; 2-tailed t-test.
